# Structural Basis of Tubulin Recruitment and Assembly by Tumor Overexpressed Gene (TOG) domain array Microtubule Polymerases

**DOI:** 10.1101/340026

**Authors:** Stanley Nithianantham, Brian D Cook, Fred Chang, Jawdat Al-Bassam

**Affiliations:** Molecular Cellular Biology Department, University of California, Davis, CA.; Department of Cell and Tissue Biology University of California, San Francisco, CA.

## Abstract

XMAP215/Stu2/Alp14 proteins accelerate microtubule plus-end polymerization by recruiting tubulins via arrays of Tumor Overexpressed Gene (TOG) domains. The underlying mechanism of these arrays as microtubule polymerases remains unknown. Here, we describe the biochemical and structural basis for TOG domain arrays in recruiting and polymerizing tubulins. Alp14 binds four tubulins via dimeric TOG1-TOG2 arrays, each with distinct exchange rates. X-ray structures reveal pseudo-dimeric square-shaped assemblies in which four TOG domains position four unpolymerized tubulins in a polarized wheel-like configuration. Crosslinking confirms square assemblies form in solution, and inactivation of their interfaces destabilizes square organizations without influencing tubulin binding. Using an approach to modulate tubulin polymerization, we determined a X-ray structure showing an unfurled assembly in which TOG1 and TOG2 uniquely bind two polymerized tubulins. Our findings suggest a new microtubule polymerase model in which TOG arrays recruit tubulins by forming square assemblies, which then unfurl facilitating their concerted polymerization into protofilaments.

## Introduction

Microtubules (MTs) are highly dynamic polarized polymers that perform critical cellular functions including forming bipolar mitotic spindles, intracellular organization, and modulating cell development and cell migration-(Akhmanova and Steinmetz, 2008, 2015). MTs are assembled from αβ-tubulin heterodimers (αβ-tubulin) and their polymerization exhibits dynamic instability arising from guanosine triphosphate (GTP) hydrolysis in β-tubulins at MT ends. However, the conformational changes promoting soluble αβ-tubulins to polymerize at MT ends remain poorly understood. The polymerization of αβ-tubulin and its GTP hydrolysis are regulated by conserved proteins that bind MT plus-ends or along MT lattices (Akhmanova and Steinmetz, 2008, 2011, 2015; Al-Bassam and Chang, 2011; Al-Bassam et al., 2010; Brouhard and Rice, 2014). The XMAP215/Stu2/Alp14 family of MT polymerases is among the best-studied MT regulators. They localize at the extreme tips of MT plus-ends and accelerate αβ-tubulin polymerization in eukaryotes (Akhmanova and Steinmetz, 2011, 2015; Al-Bassam and Chang, 2011; Maurer et al., 2014). Loss or depletion of MT polymerases is lethal in several organisms as it severely decreases MT polymerization rates during interphase resulting in shortened mitotic spindles in most eukaryotes studied (Al-Bassam et al., 2012; Cullen et al., 1999; Wang and Huffaker, 1997). MT polymerases also bind kinetochores where they accelerate MT dynamics and regulate kinetochore-MT attachment (Miller et al., 2016; Tanaka et al., 2005). MT polymerases recruit αβ-tubulins via an array of conserved Tumor Overexpressed Gene (TOG) domains (termed TOG arrays, from herein), which are critical for their function (Reber et al., 2013; Widlund et al., 2011). Arrays of TOG-like domains are conserved in two other classes of MT regulators, CLASP and Crescerin/CHE-12 protein families (Al-Bassam and Chang, 2011; Al-Bassam et al., 2010; Das et al., 2015), suggesting that arrays of TOG domains uniquely evolved to regulate diverse MT polymerization functions through binding αβ-tubulins in different intracellular settings.

Yeast MT polymerases, such as S. *cerevisiae* Stu2p and *S. pombe* Alp14, are homodimers consisting of two unique and consecutive TOG domains, TOG1 and TOG2, per subunit numbered based on their location from the N-terminus. In contrast, metazoan orthologs, such as XMAP215 and ch-TOG, are monomers including five tandem TOG domains, TOG1 through TOG5, (Al-Bassam and Chang, 2011; Brouhard and Rice, 2014). Phylogenetic analyses suggest that TOG1 and TOG2 domains are evolutionarily distinct (Al-Bassam and Chang, 2011), and the TOG3 and TOG4 domains in metazoans are evolutionarily and structurally related to the TOG1 and TOG2 domains, respectively (Brouhard et al., 2008; Fox et al., 2014; Howard et al., 2015). Thus, despite differences in TOG array organization in yeast and metazoan proteins, both groups contain at least two sets of tandem TOG1-TOG2 domains.

Structural studies contribute to our understanding of the molecular basis of TOG domain function in recruiting soluble αβ-tubulin. Each TOG domain is composed of six α-helical HEAT (Huntingtin, EF3A, ATM, and TOR) repeats, which forms a conserved paddle-shaped structure (Al-Bassam and Chang, 2011; Al-Bassam et al., 2007; Al-Bassam et al., 2006; Brouhard and Rice, 2014; Slep and Vale, 2007). X-ray structures of isolated TOG1 and TOG2 domains in complex with αβ-tubulins reveal that these domains recognize the curved αβ-tubulin conformations via inter-helical loops positioned along an edge of these paddle-shaped domains (Ayaz et al., 2014; Ayaz et al., 2012). Straightening of the curved soluble αβ-tubulins upon polymerization into MTs likely dissociates TOG domains. Our previous studies indicate that native TOG arrays from yeast or metazoan MT polymerases assemble into discrete particles upon binding αβ-tubulin (Al-Bassam and Chang, 2011; Al-Bassam et al., 2006; Brouhard and Rice, 2014). Both TOG1 and TOG2 domains are critical for MT polymerase function and their inactivation in fission or budding yeast MT polymerases results in MT function defects (Al-Bassam et al., 2012; Al-Bassam et al., 2006; Ayaz et al., 2014). Two models were proposed to explain how arrays of TOG domains function as MT polymerases: One model, which is based on studies of native TOG arrays, indicates that TOG arrays may form ordered assemblies upon binding αβ-tubulins (Al-Bassam et al., 2006; Brouhard et al., 2008). A second model, based on studies of isolated TOG domains or short TOG arrays suggests these arrays form flexible assemblies in which TOG1 and TOG2 independently recruit multiple αβ-tubulins to MT plus-ends (Al-Bassam and Chang, 2011; Ayaz et al., 2014). Distinguishing between these models requires understanding the high-resolution organization of native TOG arrays in complex with αβ-tubulin and their transitions during the αβ-tubulin recruitment and polymerization phases.

Here, we describe biochemical and structural analyses of TOG arrays during αβ-tubulin recruitment and polymerization states. We show that dimeric yeast MT polymerases recruit αβ-tubulins using TOG1 and TOG2 domains, which bind and release αβ-tubulins with unique exchange rates. X-ray structures of αβ-tubulin-TOG array complexes reveal pseudo-dimeric TOG1-TOG2 subunits form a head-to-tail square-shaped assembly, which orients αβ-tubulins in a polarized configuration. Crosslinking and mass spectrometry show dimeric yeast TOG arrays that form square conformations, confirming that these states exist in solution. Mutants in which these interfaces are inactivated show disrupted square organization, without defects in αβ-tubulin binding. Using a novel approach to promote the limited polymerization of αβ-tubulin while bound to TOG arrays, we determined a x-ray structure of an “unfurled” assembly revealing TOG1-TOG2 domains are bound onto two αβ-tubulins polymerized head-to-tail into a protofilament. These studies establish a new “polarized unfurling” model for TOG arrays as MT polymerases. In the accompanying manuscript, we present *in vitro* reconstitution studies and *in vivo* functional studies in the fission yeast *S. pombe* using two classes of Alp14 structure-based mutants, which provide evidence supporting various facets of this new model.

## Results

### Arrays of TOG1 and TOG2 domains recruit multiple αβ-tubulins via and exhibit unique affinities and exchange rates

First, we studied the binding affinities and stoichiometries of near native monomeric and dimeric yeast TOG arrays to bind αβ-tubulin. A yeast TOG array consists of TOG1 and TOG2 domains linked by 55 to 80-residue linker and can be dimerized via a C-terminal coiled-coil. Individual TOG domains bind αβ-tubulins via a narrow interface, which involves mostly ionic contacts (Ayaz et al., 2014; Ayaz et al., 2012). Thus, we explored how changes in ionic conditions influence αβ-tubulin binding stoichiometry of monomeric *S. pombe* Alp14 TOG1-TOG2 array (termed Alp14-monomer from herein, and includes residues 1-510) or dimeric Alp14 TOG array (termed wt-Alp14-dimer from herein: residues 1-690), using size-exclusion chromatography (SEC) and multi-angle light scattering (SEC-MALS) (Figure 1A, B, F; Figure 1-Supplement 1A, B, D, E and Figure 1 Supplement 2G-I) (Al-Bassam et al., 2012). At 80 or 100 mM KCl conditions, 1 μM wt-Alp14-dimer bound to four αβ-tubulins via its four TOG domains, whereas the 1 μM wt-Alp14-monomer only bound to two αβ-tubulins via two TOG domains, suggesting TOG1 and TOG2 independently recruit αβ-tubulins (Figure 1A, B, F; Figure 1-Supplement 1A-C, D-F and Figure 1 Supplement 2G-I; Table S1–S2). At 200 mM KCl conditions, however, both 1 μM wt-Alp14-monomer and wt-Alp14-dimer bound roughly half the expected αβ-tubulin compared to that at 80 or 100 mM KCl, resulting in lower-mass complexes. The latter suggests that either TOG1 or TOG2 domain, but not both, maintains binding to αβ-tubulin at 200 mM KCl conditions (Figure 1A, B, F; Figure 1 Supplement1A-C, D-F and Figure 1 Supplement 2H-I; Table S1–S2); the values for binding stoichiometry at these 100 or 200 mM ionic conditions resolve the discrepancies between our previously reported binding stoichiometries and those reported by other groups (Al-Bassam et al., 2012; Al-Bassam et al., 2006). To determine if either TOG1 or TOG2 retains αβ-tubulin binding at 200 mM KCl, we studied Alp14 mutants in which TOG1 or TOG2 αβ-tubulin binding interfaces were inactivated through mutating conserved intra-HEAT repeat turn residues (see materials and methods; Figure 1C,D; Figure 1 Supplement 1G-L). 1 μM wt-Alp14-dimer with inactivated TOG1 (termed TOG1M), which includes only active TOG2, bound αβ-tubulins at 100 mM KCl, but dissociated from αβ-tubulin at 200 mM KCl, as evidenced by poor co-migration of Alp14 with αβ-tubulin on SEC, and with the majority of αβ-tubulin migrating into a separate SEC peak (Figure 1C; Figure 1 Supplement 1G-I, Figure 1 Supplement 2J). In contrast, Alp14-dimer with an inactivated TOG2 (TOG2M), which only includes active TOG1, bound the majority of αβ-tubulin in both 100 and 200 mM KCl conditions (Figure 1D; Figure 1 Supplement 1J-L, Figure 1-Supplement 2J; Table S1–S2). Moreover, the αβ-tubulin binding molar ratios and the measured stoichiometry of TOG1M and TOG2M at 80-100 mM KCl, determined by SEC and SEC-MALS, respectively, were roughly half those measured for wt-Alp14-dimer, supporting the independent, and unique activities of TOG1 and TOG2 in recruiting αβ-tubulins (Figure 1E).

**Figure 1.**
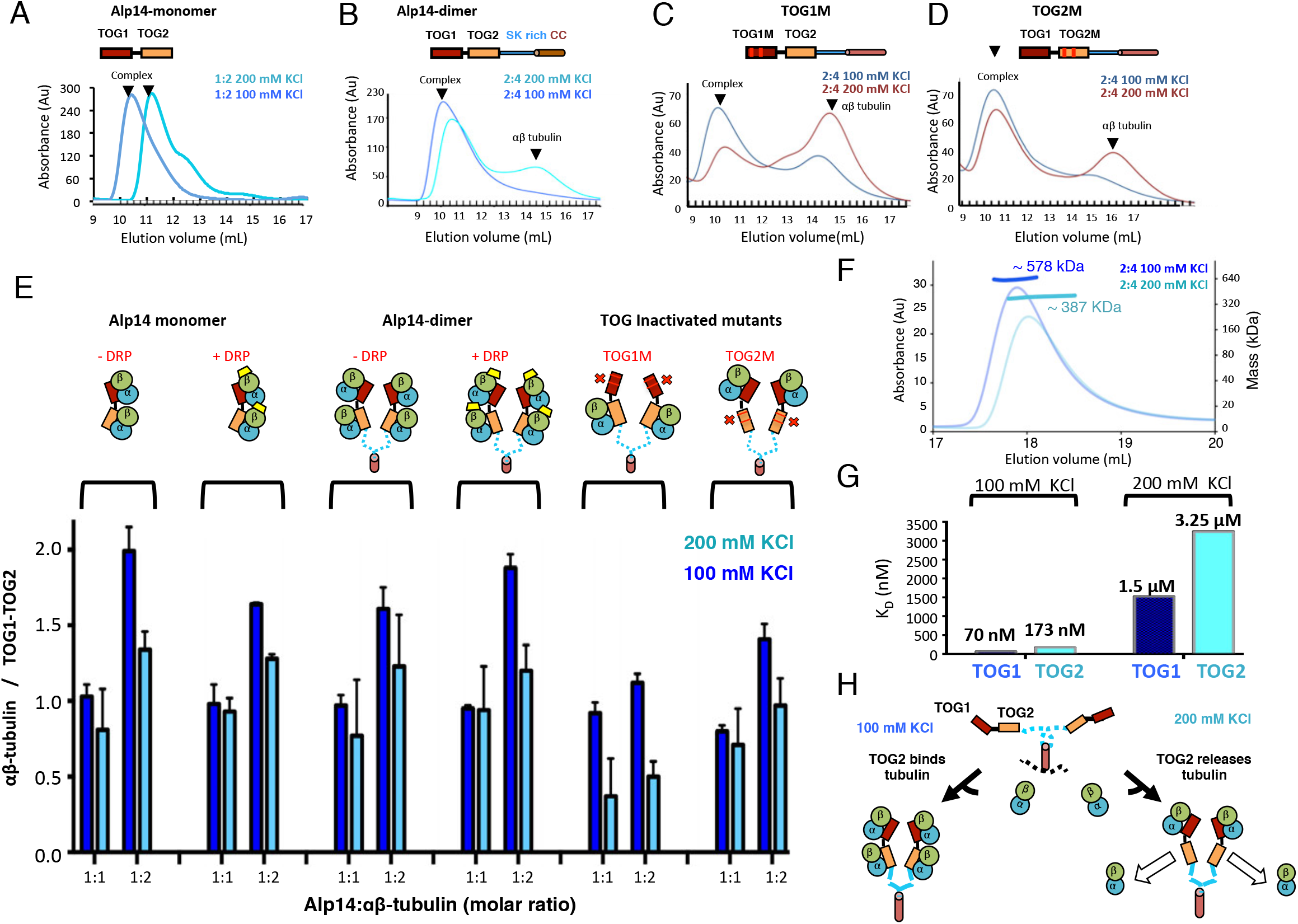
TOG1 and TOG2 domains in yeast MT polymerases both bind αβ-tubulins but exchange them at different rates. (**A-D**) Top, domain organization of the monomeric yeast Alp14 (A: Alp14-monomer), dimeric Alp14 (**B**: Alp14-dimer) and Alp14 inactivated TOG1 (**C**: TOG1M) and Alp14 with inactivated TOG2 (**D**: TOG2M). (**A** and **B**) Bottom, SEC of wt-Alp14-monomer or wt-Alp14-dimer proteins in αβ-tubulin complex at 1:2 or 2:4 stoichiometry at 100 mM and 200 mM KCl, revealing the dissociation of a substantial amount of αβ-tubulin upon increase of ionic strength. (**C** and **D**) Bottom SEC of TOG1M shows substantial αβ-tubulin dissociation at 200 mM KCl compared to Alp14-dimer-TOG2M, which releases less αβ-tubulin. (**E**) SEC-based titration measured through analyses of masses from SDS-PAGE (Table S2; materials and methods) of Alp14-monomer, Alp14-dimer, TOG1M and TOG2M with and without DRP binding, reveal the activities of TOG1 and TOG2 domains, and their nonequivalence in αβ-tubulin exchange at 200 mM KCl. Details are described in Figure 1 Supplement 1. (**F**) SEC-MALS traces for Alp14-dimer binding αβ-tubulin at 2:4 molar ratio at 100 mM KCl or 200 mM KCl, respectively. (**G**) Isothermal titration calorimetery (ITC) reveals TOG1 and TOG2 exchange αβ-tubulin with non-equivalent rates at 200 mM KCl. At 100 mM KCl, TOG1 and TOG2 both slowly dissociate from αβ-tubulin with K_D_=70 nM and K_D_=173 nM, respectively. At 200 mM KCl, TOG1 dissociates from αβ-tubulin slowly with a K_D_ =1.50 μM, while TOG2 mostly dissociates from αβ-tubulin with a K_D_=3.25 μM. ITC binding curves are shown in Figure 1 Supplement 3. (H) Model for non-equivalent activities of TOG1 and TOG2 within TOG array for recruiting αβ-tubulins.

We, next, quantitatively measured the absolute TOG1 and TOG2 affinities for αβ-tubulin and how those change due to changes in ionic strength (100-200 mM KCl) using isothermal titration calorimetery (ITC). ITC data show isolated TOG1 (residues 1-270) and TOG2 (residues 320-510) bind αβ-tubulins with roughly 2.5-fold difference in dissociation constants (Figure 1G; Figure 1-Supplement 3). At 100 mM KCl, we measured dissociation constants for TOG1 and TOG2 to be 70 and 173 nM, respectively. These data suggest both TOG1 and TOG2 exhibit a fairly high affinity for αβ-tubulin with 2.5-fold affinity difference and are nearly identical to those previously reported (Ayaz et al., 2014; Ayaz et al., 2012). However, at 200 mM KCl, we measured TOG1 and TOG2 αβ-tubulin dissociation constants at 1.5 μM and 3.2 μM, respectively, which are 20-fold lower in absolute affinity compared to those measured at 100 mM KCl. Together, our studies suggest that at 100 mM KCl or below, each TOG1-TOG2 array subunit tightly binds two αβ-tubulins, whereas at 200 mM KCl it binds one αβ-tubulin tightly via TOG1 and exchanges a second αβ-tubulin rapidly via TOG2 (Figure 1H). In the accompanying manuscript, we describe the implications of this ionic strength change in modulating the MT polymerase activity through influencing TOG2 (Cook et al., 2018). We also present *in vitro* and *in vivo* studies for TOG1M and TOG2M, suggesting that TOG1 and TOG2 domains, with their unique exchange rates for αβ-tubulin serve unique and non-additive roles in MT-plus end tracking and MT polymerase activity, respectively (Cook et al., 2018).

### TOG arrays form a pseudo-dimeric square assembly that orients αβ-tubulins

Our biochemical studies suggest that structural studies of TOG arrays:αβ-tubulin complexes must be conducted at 80-100 mM KCl and at high concentrations to avoid αβ-tubulin dissociation from TOG2 domains. To increase complex homogeneity and inhibit αβ-tubulin self-assembly under such conditions, we utilized designer ankyrin repeat protein, Darpin-D1 (termed DRP from herein), which specifically binds β-tubulin’s polymer-forming interface (Pecqueur et al., 2012). We studied if DRP binding to αβ-tubulin influences the ability of wt-Alp14-monomer or wt-Alp14-dimer to bind αβ-tubulins. Binding molar ratios and stoichiometries were measured for DRP bound-αβ-tubulin wt-Alp14 complexes by SEC and SEC-MALS at 80-100 mM KCl conditions (Figure 1E; Figure 1 Supplement 2A-G; Table S1–S2). These studies suggest that DRP does not affect the simultaneous binding of multiple αβ-tubulins to TOG arrays in monomer or dimer forms. At 1 μM concentration, wt-Alp14-dimer formed a complex with four αβ-tubulin and four DRP at a molar ratio of 2:4:4 (Figure 1E; Figure 1 Supplement 2D-F). The SEC-MALS measured wt-Alp14-monomer:αβ-tubulin:DRP complex is 1:2:2 stoichiometry indicating each TOG1-TOG2 subunit binds two αβ-tubulins each of which binds a DRP (Figure 1E; Figure 1 Supplement 2A-C, G; Table S1). These ability of these tubulin to bind stoichiometric amounts of DRP suggest that αβ-tubulins recruited by TOG array are in a non-polymerized state upon their initial association. This feature is consistent with a reported lack of cooperativity described between TOG1 and TOG2 in binding to αβ-tubulins (Ayaz et al., 2014). Accordingly, we used this strategy to identify crystallization conditions using yeast MT polymerase orthologs from a variety of organisms (see materials and methods).

Crystals of the *Saccharomyces kluyveri* ortholog of Alp14 (termed sk-Alp14-monomer; residues 1-550) bound to αβ-tubulins and DRP grew in conditions similar to those used for SEC and SEC-MALS (Figure 2 Supplement 1A). Using crystals with either a native sk-Alp14-monomer or a sk-Alp14-monomer with a modified TOG1-TOG2 linker sequence, termed sk-Alp14-monomer-SL (see materials and methods; Figure 2 Supplement 2), we determined X-ray structures for 1:2:2 sk-Alp14 TOG1-TOG2: αβ-tubulin: DRP from complexes using sk-Alp14-monomer and sk-Alp14-monomer-SL with molecular replacement (see materials and methods) at 4.4 and 3.6-Å resolution, respectively (Table S3 and Figure 2 Supplement 1B, C). In the structures, TOG1 domains were clearly differentiated from TOG2 domains by their conserved C-terminal extension and jutting α-helix, that were unambiguously identified in density-modified maps (Figure 2 Supplement 1D, E). Each asymmetric unit contained two wheel-shaped assemblies (Figure 2 Supplement 1F), representing two sets of alternating TOG1 and TOG2 domains oriented in a square-like conformation (termed the TOG square), with each TOG domain binding a DRP capped αβ-tubulin on its outer edge. Excluding the 10-residues TOG1-TOG2 linker region immediately preceding TOG2, the remaining 40 residues of the linker were disordered (Figure 2-Supplement 1F-I).

The dimension of each wheel-like assembly is 210 × 198 × 60-Å (Figure 2A). The 2:4:4 stoichiometry observed in the X-ray structure matched the stoichiometry measured for wt-Alp14-dimer:αβ-tubulin: DRP complexes (Figure 1B, F; Figure 1 Supplement 2D-F). We anticipate sk-Alp14-monomer formed dimeric organization, despite a lack of dimerization domains, due to their high concentration during crystallization. The X-ray structures revealed two TOG1-TOG2 subunits in a pseudo-dimeric assembly forming the core of these complexes. In a TOG square, each TOG domain is bound to a curved αβ-tubulin capped by a DRP through its outward-facing binding interface, and is minimally contacted by the neighboring TOG-bound αβ-tubulin (Figure 2A; Figure 2 Supplement 1F-H). The distances and interaction patterns between residues of α-tubulin and DRP bound onto a neighboring β-tubulin indicate that DRP only interacts with its cognate β-tubulin does not bind a neighboring α-tubulin (Figure 2 Supplement 2H-K). The latter suggests DRP has no effect on stabilizing each TOG square assembly. DRP binding rather only caps β-tubulin, presenting a significant impediment to the polymerization of αβ-tubulin while bound to the TOG array. The αβ-tubulins bound onto the TOG square are positioned in a polarized orientation, by virtue of the asymmetry in the TOG domain αβ-tubulin interface and pseudo-dimeric TOG1-TOG2 subunit interfaces within the TOG square (see below). The β-tubulin on a TOG1 bound αβ-tubulin is rotated roughly 90° from its polymer-forming interface relative to the adjacent α-tubulin on a TOG2 bound αβ-tubulin (Figure 2B).

**Figure 2:**
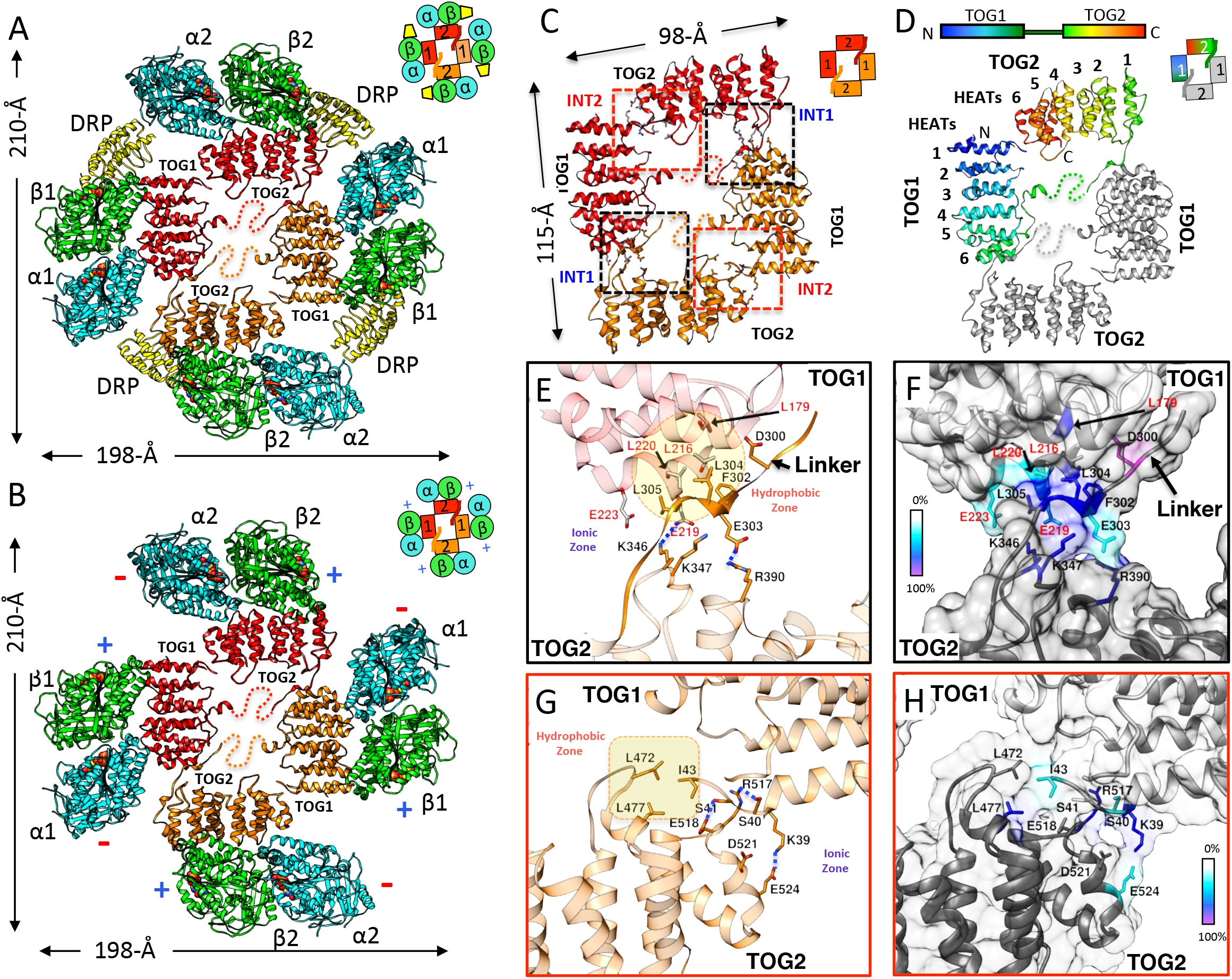
X-ray Structures reveal αβ-tubulins bound in a wheel-like organization around a pseudo-dimeric TOG square complex. (**A-B**) 3.6-Å X-ray crystal structure of the *S. kluyveri* 1:2:2 sk-Alp14:αβ-tubulin: DRP reveals pseudo-dimeric head-to-tail subunits (red and orange) in a TOG square assembly consisting of four TOG domains bound to four αβ-tubulins in a wheellike (α-tubulin shown in cyan and β-tubulin shown in green) organization. (**A**) It shows structure with DRP (yellow) bound to each αβ-tubulin. (**B**) It shows structure with DRP computationally removed. Each αβ-tubulin (α1β1) is positioned 90° rotated from its polymer-forming interface on its neighboring αβ-tubulin (α2β2). (**C**) Pseudo-dimeric TOG1-TOG2 subunits, shown in orange and red, respectively, form a head-to-tail TOG square (inset). Interface 1 is formed by N-terminus of TOG2 and the TOG1-TOG2 linker binding to the C-terminus of the TOG1 domain of a second subunit, forming a 90° corner. Interface 2 is formed by the N-terminus of TOG1 binding the C-terminus of TOG2 within the same subunit in a 90° corner (Figure 2 Supplement 11). (**D**) Rainbow view of TOG1-TOG2 with N- and C-termini displayed in a blue-to-red color gradient, while the other subunit is displayed in grey. Each TOG is composed of six HEAT repeats (numbered). (**E**) Close-up view of interface 1. A hydrophobic zone stabilizes interface1 (yellow and highlighted by red outline) involving Leu220 (L220) and Leu217 (L217) of the TOG1 inter-HEAT 5-6 loop, Leu179 (L179) of the HEAT 6 A-helix in TOG1 (red ribbon) stabilized by linker residues (solid orange) Phe302 (F302), Leu304 (L304), and Leu305 (L305). An ionic zone guides interface 1 involving Glu219 (E219) of TOG1 inter-HEAT 5-6 loop and Glu305 (E305) of the TOG1-TOG2 linker, forming salt bridges with Lys346 (K346) and Lys347 (K347) of the TOG2 (light orange) inter-HEAT 1-2 loop and Arg390 (R390) of the TOG2 HEAT 2,3 loop, respectively. (**F**) Close-up view of interface 1, as in C, displaying residue conservation based on the alignment shown in Figure 2-Supplement 2. (G) Close-up view of interface 2. A hydrophobic zone stabilizes interface 1 involving Leu477 (L477) and Leu472 (L472) of the TOG2 inter-HEAT4-5 loop with Ile43 of the TOG1 inter-HEAT1-2 loop. Ionic zone selectively guides interface 2, involving Lys39 (K39) and Ser41 (S41) of the TOG1 inter-HEAT1-2 loop and helix 1B with Arg517 (R517), Glu518 (E518), Asp521 (D521), and Glu524 (E524) of the TOG2 inter-HEAT5-6 loop and A-helix. (**H**) Close-up view of interface 2, as in D, displaying reside conservation based on the alignment in Figure 2 Supplement 2.

### Two interfaces stabilize pseudo-dimeric TOG array into a TOG square

The X-ray structures reveal each TOG square is a dimer of TOG1-TOG2 array subunits assembled head-to-tail from alternating TOG1 and TOG2 domains. TOG domains are aligned along their narrow edges, analogous to four links attached head-to-tail forming an asymmetric square-like complex with two edges slightly longer than their orthogonal edges (Figure 2C, D). Two contact sites, which we term interfaces 1 and 2, stabilize the TOG square. These interfaces are formed by interactions formed via inter-HEAT repeat loops of each TOG domain, which are located on the opposite edges from the αβ-tubulin-binding sites. Although TOG1 and TOG2 domains are each 60-Å long, the TOG square assembly is slightly rectangular with 115-Å by 98-Å dimensions due to wider overlaps between TOG1 and TOG2 domains leading to 10-Å stagger at interface 1 sites, in contrast to a direct end-on corner-like interface 2 sites. Both interfaces 1 and 2 are stabilized by hydrophobic packing and ionic interaction zones (Figure 2E-H). Interface 1 packs a 668-Å^2^ surface area, and positions the TOG1 C-terminus at 90° to a 10-residue segment of the TOG1-TOG2 linker and the N-terminus of TOG2. The TOG1-TOG2 linker sequence forms an extended polypeptide that critically bridges interactions between TOG1 HEAT repeat 6 α-helix/inter-HEAT 5-6 loop segment and the TOG2 inter-HEAT repeat 1-2/2-3 loop segments (Figure 2E, F). Interface 2 packs a 290-Å^2^ surface area and positions the TOG2 C-terminus at 90° to the N-terminus of TOG1 (Figure 2G, H) In interface 2, the TOG2 inter-HEAT repeat 4-5 loop interacts with the TOG1 inter-HEAT repeat 1-2/HEAT1 α--helix (Figure 2G). The residues forming interface 1 and interface 2 within TOG1, TOG2 and linker regions are either moderately or highly conserved (Figure 2F, H; Figure 2 Supplement 2). The total buried surface area stabilizing two sets of interfaces 1 and 2 in a TOG square is 1930 Å^2^, which is moderate in size and dispersed for such a large assembly. This suggests that this conformation maybe meta-stable and that DRP binding and its inhibition of αβ-tubulin polymerization may have stabilized this intermediate. ‘

### Dimeric Alp14 TOG arrays forms a TOG square conformation in solution

Next we examined and chemically trapped the direct physical interactions between TOG1 and TOG2 based on the interfaces observed in TOG square structure using cysteine crosslinking. We generated mutants with specific cysteine pairs within the two sides of interface 1 (S180C, L304C) or interface 2 (S41C and E518C) in native dimeric sk-Alp14 (termed sk-Alp14-dimer; residues 1-724) (Figure 3A, B). We tested whether these interfaces form inter-subunit contacts in dimeric TOG array by crosslinking via disulfide oxidation. A 110-kDa crosslinked species was observed in all conditions where soluble αβ-tubulin was added and mass spectrometry (LC/MS-MS) confirmed this intermediate is indeed a crosslinked α and β-tubulin heterodimer (Figure 3C, D; Figure 3 Supplement 1A). We also observed a ∼170-kDa species which specifically formed in the sk-Alp14 S180C-L304C mutant, and not in the native sk-Alp14-dimer or the sk-Alp14-S41C-E518C mutants. Furthermore, this 170-kDa intermediate was also not observed with sk-Alp14-S180C-L304C without αβ-tubulin (Figure 3D). Mass-spectrometry (LC/MS-MS) confirmed this 170-kDa intermediate is indeed the sk-Alp14-S180C-L304C protein. Next, we mapped the cysteine residues involved disulfides in sk-Alp14-S180C-L304C mutant through peptide disulfide mapping after differential alkylation and mass-spectrometry (see materials and methods). This approach revealed only two classes of peptides in sk-Alp14 S180C-L304C with 105-Da mass added onto cysteines, suggesting they were engaged in disulfides, with the following sequence boundaries: 297-320 and 179-189 (Figure 3 Supplement 1B). These two peptide sequences represent TOG1 inter-HEAT-repeat and TOG1-TOG2 linker regions both of which are involved in forming interface 1 in the x-ray structures (Figure 2). All the remaining peptides with cysteine residues that were identified in sk-Alp14 S180C-L304C included 57-Da in added mass, suggesting they were in the reduced form and did not form disulfides. Thus, these data directly provide independent support that interface 1 of the TOG square conformation forms in sk-Alp14-dimer in solution outside the crystallographic setting, and is indeed the inter-subunit dimeric interface between two TOG arrays subunits, as visualized in the crystal structures (Figure 3A, B).

**Figure 3:**
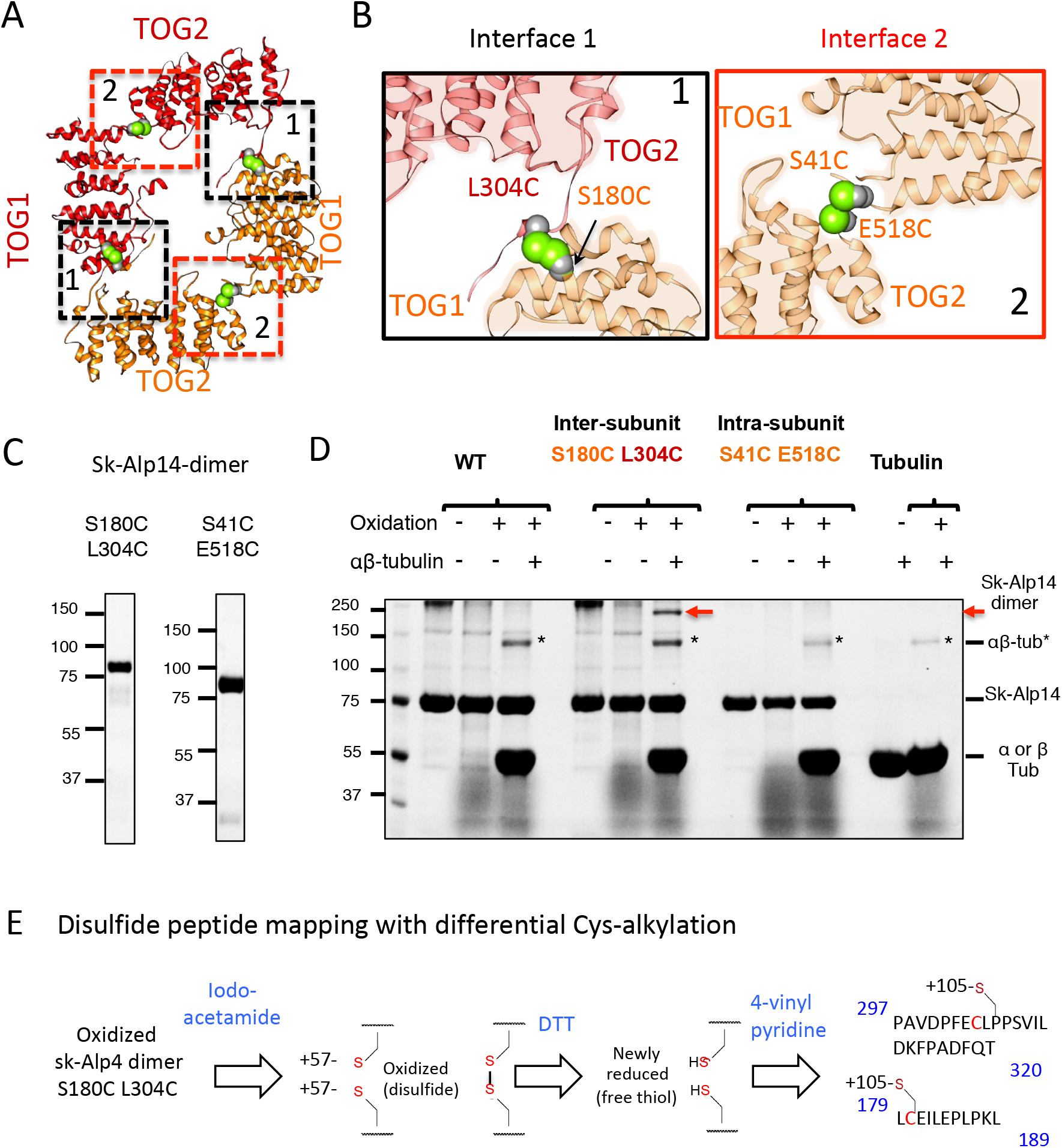
Cysteine crosslinking reveal that dimeric TOG arrays form TOG square conformations in solution. (**A**) TOG square structure showing two cysteine pairs (green space fill) mutated in interfaces 1 (black-dashed lines) and 2 (red-dashed lines). (**B**) Close-up views of interfaces 1 (left) and 2 (right) showing the S180C-L304C and E518C-S41C residue pairs, respectively (green space fill) in the sk-Alp14. (**C**) SDS-PAGE of SEC purified sk-Alp14 S180C-L304C and E518C-S41C. (**D**) Crosslinking studies of cysteine structural based mutants using oxidizing conditions and αβ-tubulin binding, as described by (+) and (-). The α β-tubulins form an intermediate in oxidizing conditions observed in all conditions that include αβ-tubulin (marked*). S180C-L304C sk-Alp14 forms a dimeric 170-kDa intermediate upon oxidization and αβ-tubulin binding (red arrow). In contrast, wt-Alp14 or E518C-S41C sk-Alp14 does not form this intermediate. (below) Disulfide peptide mapping of cysteines in sk-Alp14 S180C-L304C using differential alkylation, and LC/MS-MLS. We used mass-spectrometry (LC/MS-MS) after a differential alkylation strategy (Figure 3 Supplement 1D) to map peptides with disulfides. Briefly, oxidized sk-Alp14-S180C-L304C (170-kDa) SDS-PAGE purified bands were subjected to proteolysis, treated with iodoacetamide. Under these conditions, 57-Da mass is added onto peptides with reduced cysteines (free thiols), without affecting disulfides. Dithiothreitol was then used to reduce peptides with disulfides, and then are treated with 4-Vinyl Pyridine, which adds 105-Da in mass onto peptides with newly formed free thiols. Using LC/MS-MS, peptides with modified cysteines were identified based on masses added to cysteines. Scheme is shown and resulting sk-Alp14 peptides are shown (details provided in Figure 3 Supplement 1D).

### Inactivation of interfaces 1 and 2 disrupts TOG square assembly but does not disrupt αβ-tubulin binding

We explored the role of Interfaces 1 and 2 in formation of the TOG square assembly and the binding of TOG arrays to αβ-tubulins, We generated three Alp14-dimer mutants which harbor either partially or fully disrupted interfaces 1 and 2 sites (Figure 4A). We targeted disruption of salt bridges or hydrophobic zones in interfaces 1 and 2 by mutating conserved alanines, leucines or glutamates (Figure 4A). Charged residues were either replaced with alanines or residues of the opposite charge, and hydrophobic residues were replaced with charged residues to dissociate hydrophobic interactions. We mutated eight residues to disrupt interface 1 (termed INT1), seven residues to disrupt interface 2 (termed INT2) or fifteen residues to disrupt both interfaces 1 and 2 (termed INT12) (Figure 4A; see materials and methods). The αβ-tubulin binding activities of INT1, INT2 and INT12 were compared to wt-Alp14-dimer in using SEC and SEC-MALS. The three mutants bound a nearly identical amount of αβ-tubulin compared to wt-Alp14-dimer (Figure 4B-E; Figure 4 Supplement 1). INT1, INT2, INT12 bound roughly four αβ-tubulins at 100 mM KCl as assessed by SEC-MALS (Table S1–2; Figure 4B-E) and roughly half of the bound αβ-tubulin dissociates at 200 mM KCl as measured by SEC (Figure 4B-E; Figure 4 Supplement 1).

**Figure 4:**
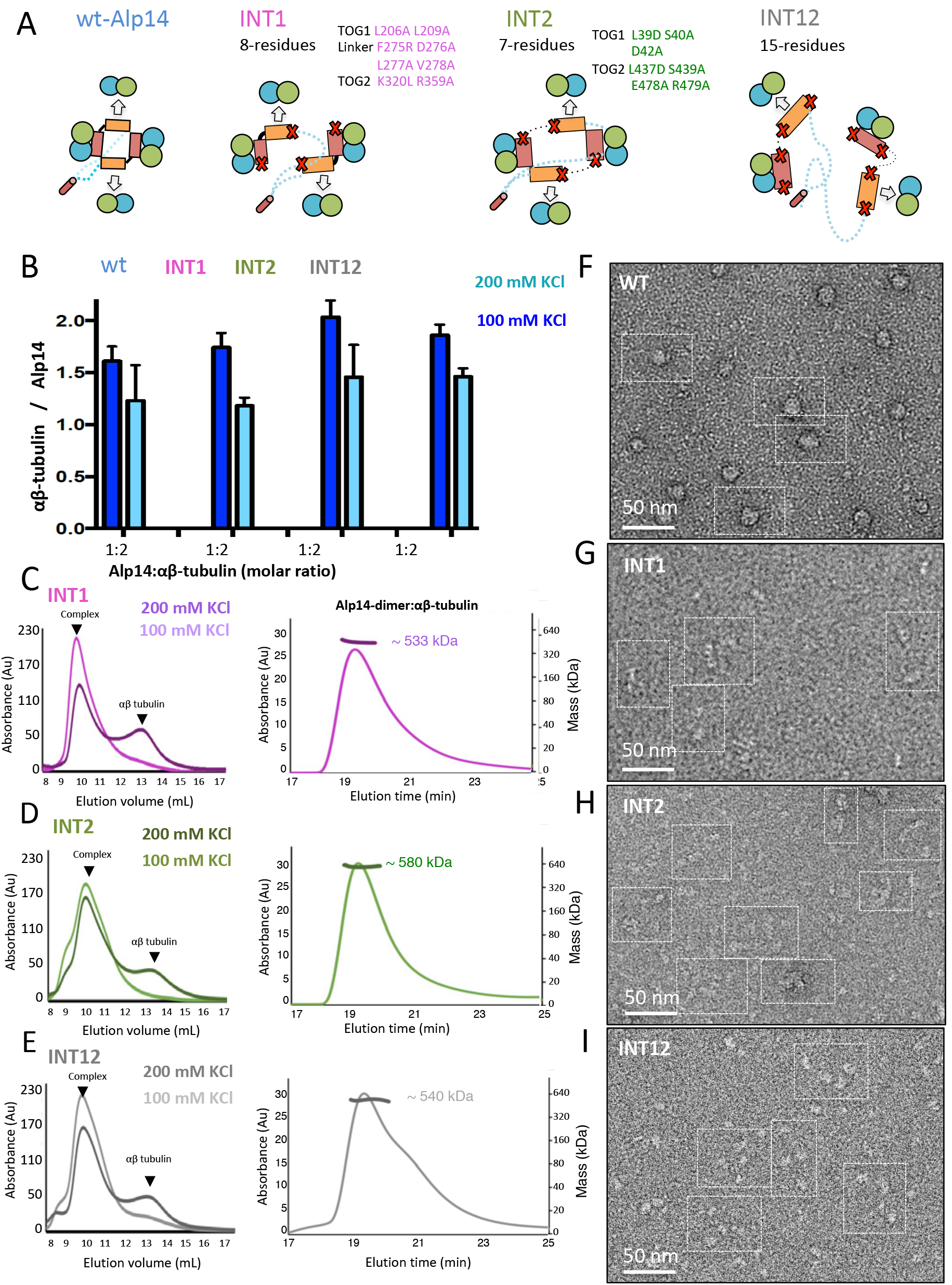
Inactivation of interfaces 1 and 2 destabilizes TOG square organization without affecting αβ-tubulin binding. (**A**) Generation of structure-based TOG square assembly defective mutants through inactivation of Interfaces 1 in the INT1 mutant (8 residue mutant), interfaces 2 in the INT2 mutant (7 residues) or both interfaces 1 and 2 in the INT12 mutant (15 residue mutant) (**B**) summary of SEC measured αβ-tubulin binding molar ratio of INT1, INT2 and INT12 compared to wt-Alp14-dimer suggests no defects αβ-tubulin binding at 100 mM KCl (blue) and a similar decrease in αβ-tubulin binding upon 200 mM KCl ionic strength increase (cyan) (**C**) Left, SEC of INT1:αβ-tubulin complexes at 2:4 stoichiometry at 100 mM KCl (light Pink) and 200 mM KCl (Dark pink), revealing the dissociation of half of the αβ-tubulin bound upon increase of ionic strength. Right, SEC-MALs of INT:αβ-tubulin complex at 100 mM KCl reveal a mass of 570 KDa. (**D**) Left, SEC for INT2:αβ-tubulin complex at 100 mM KCl (light green) and 200 mM KCl (Dark Green), as described in C. Right, SEC-MALS for INT2:αβ-tubulin complex. (**E**) Left, SEC for INT12:αβ-tubulin complex at 100 mM KCl (Light Grey) and 200 mM KCl (Dark Grey) as described in C. Right, SEC-MALS for INT12:αβ-tubulin complex. (**F**) Negative stain EM reveals wt-Alp14-dimer: αβ-tubulin complex at 100 mM KCl revealing wheel-shaped assemblies that are 15-19 nm in diameter as previously described (Al-Bassam et al 2006). (**G**) INT1 form elongated conformations with either 8 nm disconnected densities or 16 nm-minifilaments (H) INT2 forms extended necklace shaped or extended 16 nm minifilaments. (**I**) INT12 forms only necklace shaped assemblies with four interconnected 8-nm densities, which likely represent flexibly connected TOG-αβ-tubulin complexes. Additional information found in Figure 4 Supplement 1.

We next studied the conformation of the four αβ-tubulin bound wt-Alp14-dimer complex using negative stain EM and compared those to the αβ-tubulin complexes of either the INT1, INT2 or INT12 mutants using negative stain EM at 80 mM KCl. The 4:2 wt-Alp14-dimer:αβ-tubulin complexes form 15-19 nm diameter compact circular complexes, the overall similar dimensions to similar to x-ray structure (Figure 2) and match the general features we previously described for yeast Stu2p-tubulin or xmap215-tubulin (Al-Bassam et al., 2006; Brouhard et al., 2008). The heterogeneity of these negative stain images due to tubulin dissociation prevented further image analyses of these data. The 4:2 INT1 complexes form open assemblies with either 8 nm long densities or 16-nm long short filament assemblies. This suggests that in the INT1-αβ-tubulin complex, the TOG1-TOG2 arrays do not form compact square dimer and that each TOG1-TOG2 subunit would bind an αβ-tubulin forming an 8 nm density. The TOG1-TOG2 array may bind two αβ-tubulins polymerized into 16-nm filaments. INT12:αβ-tubulin complexes show 8-nm size “beads in a necklace” conformation (Figure 4I), suggesting disorganized TOG1-TOG2 array connected by flexible linkers. INT12-αβ-tubulin complex shows the most disorganized state when compared to wt-Alp14-dimer, INT1 and INT2. The INT12-αβ-tubulin complex clearly consists of four 8-nm densities likely representing the four flexibly connected TOG-bound αβ-tubulin. Our biochemical and negative stain-EM studies suggest that INT1, INT2 and INT12 all bind four αβ-tubulins, similar to wt-Alp14-dimer but their conformations are disorganized due the destabilization of interface 1 and 2. The square TOG array organization is clearly disrupted in these mutants without any effect on αβ-tubulin binding via TOG1 and TOG2 domains. INT1-αβ-tubulin shows the propensity to form two polymerized αβ-tubulins, suggesting spontaneous in-solution αβ-tubulin polymerization occurs in the case of INT1-destabilization. In the accompanying manuscript (Cook et al., 2018), we describe MT polymerase and plus-end activities of the INT1, INT2 and INT12 mutants using *in vitro* reconstitution and *in vivo* functional studies in fission yeast cells, revealing defects in processive MT plus-end tracking leading to loss of MT polymerase activities suggesting the TOG square is an essential conformation of the MT polymerase cycle.

### Unfurling of TOG arrays promote concerted αβ-tubulin polymerization into protofilaments

The TOG square conformation shows how αβ-tubulins are recruited but does not reveal how TOG arrays drive αβ-tubulin polymerization. We hypothesized that the TOG square structure may undergo a subsequent conformational change to promote polymerization of the recruited αβ-tubulins. To explore this transition, we created a biochemical approach to partially release αβ-tubulin polymerization by relieving the DRP inhibition. We reasoned that a structural transition may occur more readily if the DRP dissociates from β-tubulin in a crystallization setting. We developed a strategy to conditionally release αβ-tubulin polymerization while recruited into TOG arrays by using a weakened affinity DRP. We reasoned that the increased dissociation of DRP may allow complexes to form polymerized αβ-tubulin intermediates in steady state. To accomplish this, we removed the N-terminal ankyrin repeat of DRP (termed DRPΔN from herein). We measured DRPΔN affinity using ITC revealing a three-fold decrease in its αβ-tubulin binding affinity as compared to DRP (Figure 5A, B). During purification, complexes of 1:2:2 sk-Alp14-monomer:αβ-tubulin: DRPΔN behaved similarly to those assembled with DRP on SEC (Figure 5C; Figure 5 Supplement 1). However, under identical crystallization conditions to those used to obtain the 4:4:2 TOG square conformation, we obtained crystals which exhibited a distinct rectangular morphology and using sk-Alp14-monomer or sk-Alp14-dimer αβ-tubulin: DRPΔN complexes, (see Materials and Methods; Figure 5-Supplement 2A). These crystals formed in the same space group (P2i) with distinct unit cell dimensions compared to the TOG square crystals (Table S3). These crystals exclusively formed only when DRPΔN was used with αβ-tubulin:sk-Alp14-monomer or dimer complexes. We determined an X-ray structure to 3.3-Å resolution by molecular replacement using these crystals (Figure 5-Supplement 2B). The structure revealed a novel assembly consisting of complexes with the stoichiometry of 1:2 sk-Alp14-monomer:αβ-tubulin and a single DRPΔN positioned on the top end of TOG2 bound β-tubulin (termed the 1:2:1 structure; Figure 5E, F; Figure 5-Supplement 2C, D).

**Figure 5.**
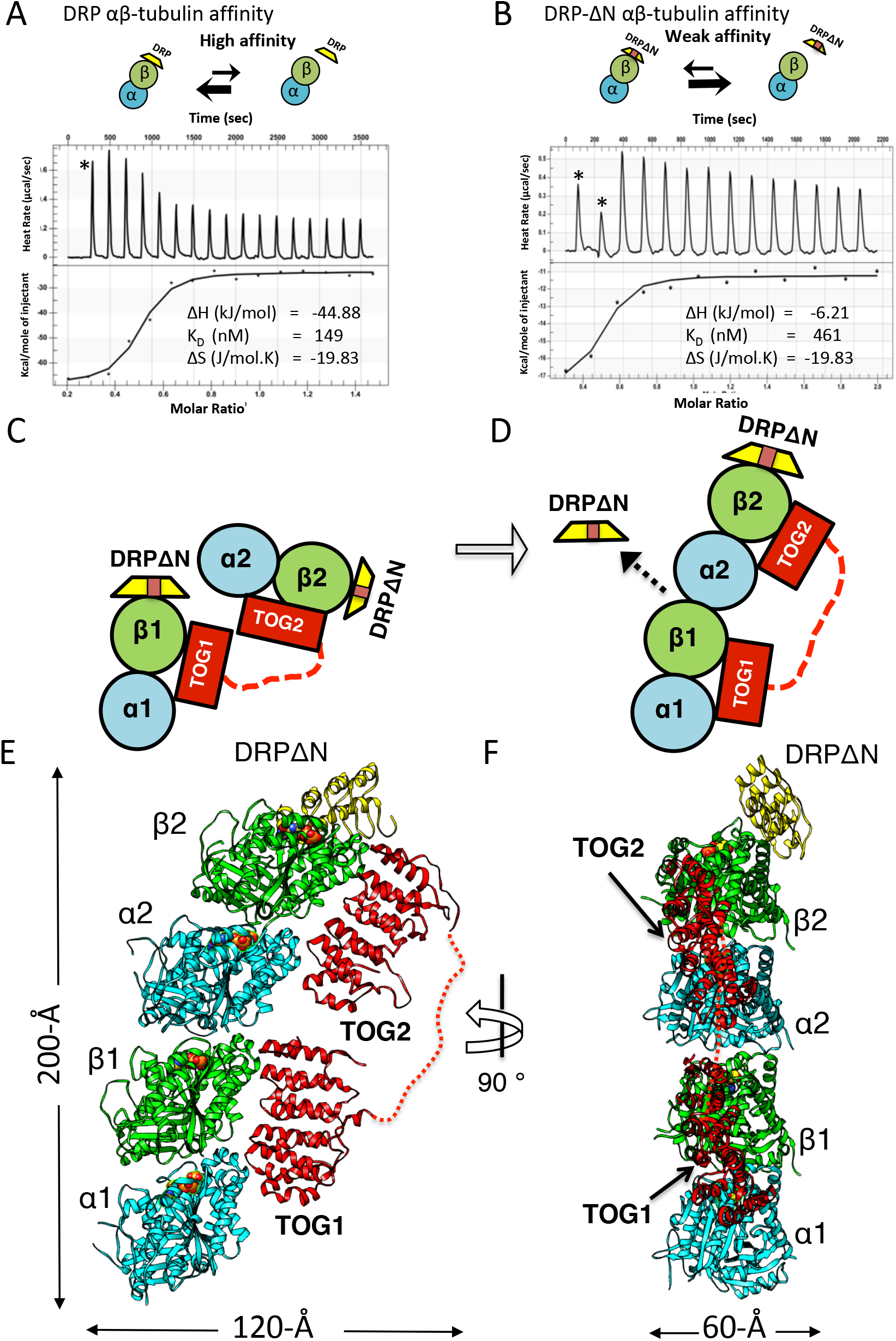
X-ray structure of 1:2:1 TOG-array:αβ-tubulin: DRPΔN reveals Unfurled TOG arrays with TOG1 and TOG2 bound to two polymerized αβ-tubulins. (**A, B**) Top, schemes DRP and DRPΔN binding to αβ-tubulin. DRP-shifts the equilibrium towards dissociation from αβ-tubulin. Bottom, Isothermal titration calorimetery studies reveal a three-fold affinity decrease in DRPΔN (461 nM) compared to DRP binding to αβ-tubulin (149 nM). (**C, D**) Two schematic views of the transition of the TOG1-TOG2 αβ-tubulin complex from the TOG square (only half is shown) to the unfurled conformation upon DRPΔN dissociation. (**E, F**) Two orthogonal views of the 3.3-Å X-ray structure of 1:2:1 sk-Alp14 (red):αβ-tubulins (cyan and green): DRPΔN (yellow) complex, indicating a polymerized protofilament state. TOG2 and TOG1 are bound to the upper (α2β2) tubulin and lower (α1β1) tubulin.

The refined 3.3-Å 1:2:1 x-ray structure reveals an extended conformation with two αβ-tubulins polymerized head-to-tail in a curved protofilament (Figure 5E, F). In this conformation, TOG1 and TOG2 do not form any interactions with each other and their adjoining linker becomes disordered (Figure 5E, F; Figure 5 Supplement 2C, D). TOG1 and TOG2 are specifically bound to the lower and upper αβ-tubulins, respectively, of a highly curved protofilament. Only a single DRPΔN caps the TOG2 bound αβ-tubulin (Figure 5E). Compared to the TOG square, this “unfurling” rearrangement requires 68°-rotation and 32-Å translation of TOG2:αβ-tubulin hinging around interface 2 and TOG1-αβ-tubulin (Figure 6A, B). This transition promotes the concerted polymerization of TOG2:αβ-tubulin onto the plus-end of TOG1:αβ-tubulin, and consequently driving the dissociation of the second DRPΔN (Figure 5C, D). The two αβ-tubulin polymer in this complex is a highly curved protofilament (16.4° inter-dimer interface), and it displays ∼3° more curvature than RB3/stathmin-αβ-tubulin curved protofilament structure (Table S5; Figure 5 Supplement 2E) (Nawrotek et al., 2011). Comparison of αβ-tubulin dimer structure (α2β2) within this 1:2:1 structure to the unpolymerized αβ-tubulin structure revealed polymerization is associated with a 5° rotation and 10-Å translation in the T7 loop and H8 helix in the TOG2-bound α-tubulin, which engages TOG1-bound β-tubulin elements and the E-site bound GDP nucleotide (Figure 6C, D). The latter conformational change likely stabilizes inter-dimer αβ-tubulin interfaces (Figure 5D), burying a 1650-Å inter-dimer interface (Figure 5E, F). This conformational change occurs at similar site to those observed during the MT lattice GTP hydrolysis transition (Alushin et al., 2014). Thus the 1:2:1 unfurled structure represents a concerted αβ-tubulin post-polymerization intermediate promoted by a TOG1-TOG2 subunit prior to straightening the protofilament.

**Figure 6:**
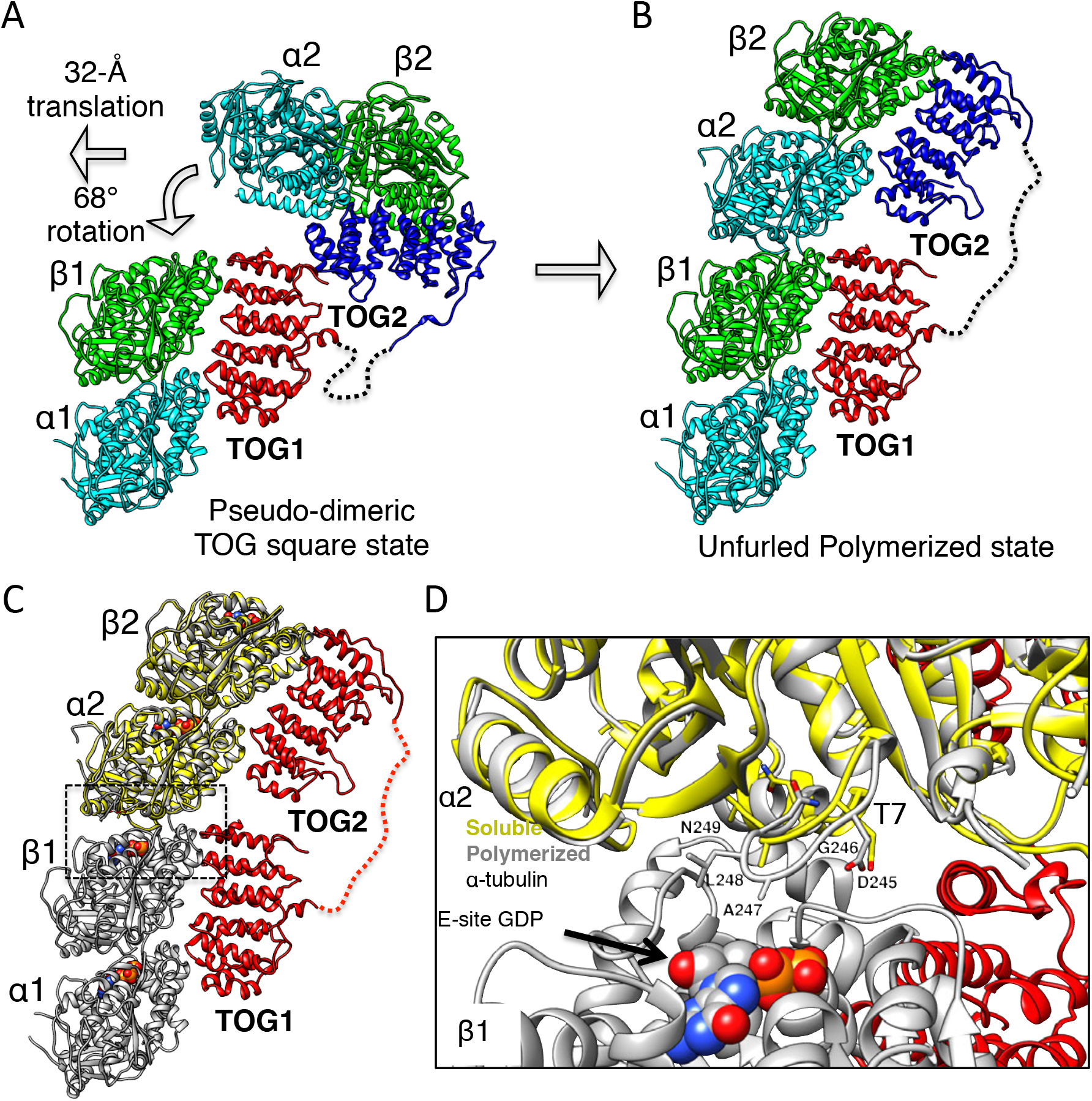
Unfurling of TOG array involves TOG2 rotation, translation around TOG1 and αβ-tubulin inter-dimer interface assembly. (**A** and **B**) Conformational change of TOG2 (blue) around TOG1 (red) while each is bound to αβ-tubulin (green and cyan) from a corner subunit in the wheel assembly (left) and in the extended conformation (right). TOG2 rotates by 32° and translates by 68 Å upon release to drive αβ-tubulin polymerization into a protofilament. (**C**) Superimposing unpolymerized αβ-tubulin (yellow) onto the α2β2-tubulin shows a conformational change in α-tubulin at the inter-dimer interface induced by polymerization. (**D**) Close-up view of the polymerized inter-dimer interface. Unpolymerized αβ-tubulin (yellow) is superimposed onto α2β2 (grey) of 1:2:1 structure. The α2-tubulin T7 loop and H8 helix engage the β1-tubulin GDP nucleotide through a T7 loop 5-Å translation and H8 helix 5° rotation involving residues Asp 245 (D245), Gly 246 (G246), Ala 247 (A247), and Leu 248 (L248).

### Modeling the interaction of the TOG square and unfurled structures with MT protofilament plus-ends

We next evaluated how x-ray structures for αβ-tubulin loaded TOG square and unfurled assemblies may dock onto protofilament tips at MT-plus ends. Atomic models of for these states were overlaid onto the terminal αβ-tubulins of the curved GTP or GDP tubulin protofilament models (Figure 7). The four αβ-tubulin bound TOG square assembly x-ray structure (Figure 2) was superimposed onto the terminal αβ-tubulin at protofilament ends in two docking orientations, either via the αβ-tubulins bound onto TOG1 or TOG2 (Figure 7A, B). We observe mild steric surface overlap between the four αβ-tubulin loaded TOG square and the protofilament when TOG1-αβ-tubulin is docked onto the protofilament end. This steric overlap caused by overlap between αβ-tubulin-TOG2 from the second TOG square subunit and the penultimate αβ-tubulin from the protofilament end (Figure 7A; Figure 7 Supplement 1A). In contrast, we observe no steric contact when the TOG square is docked via αβ-tubulin-TOG2. In this orientation, the TOG1-αβ-tubulin from the second subunit is retracted by 10-Å from the penultimate αβ-tubulin in the protofilament (Figure 7B). The differences between steric overlap between the TOG square and the protofilament in these two docking orientations are due to slightly asymmetric length and width dimensions of the TOG square, which are caused by stagger at interface 1. These differences suggest that the destabilization of the TOG squares is more likely if TOG1-αβ-tubulin docks onto the protofilament plus end, in contrast to TOG2-αβ-tubulin docking. The slow exchange rate of TOG1-αβ-tubulin makes this more likely while the rapid exchange of αβ-tubulin by TOG2 makes this less likely. The unfurled 1:2 TOG1-TOG2:αβ-tubulin assembly can only docked using TOG1-αβ-tubulin onto the protofilament plus-end, and suggests that TOG2:αβ-tubulin is positioned the furthest away from the MT-plus-end in this conformation. These models were used to assemble steps for a new MT polymerase model described in the discussion (Figure 8).

**Figure 7:**
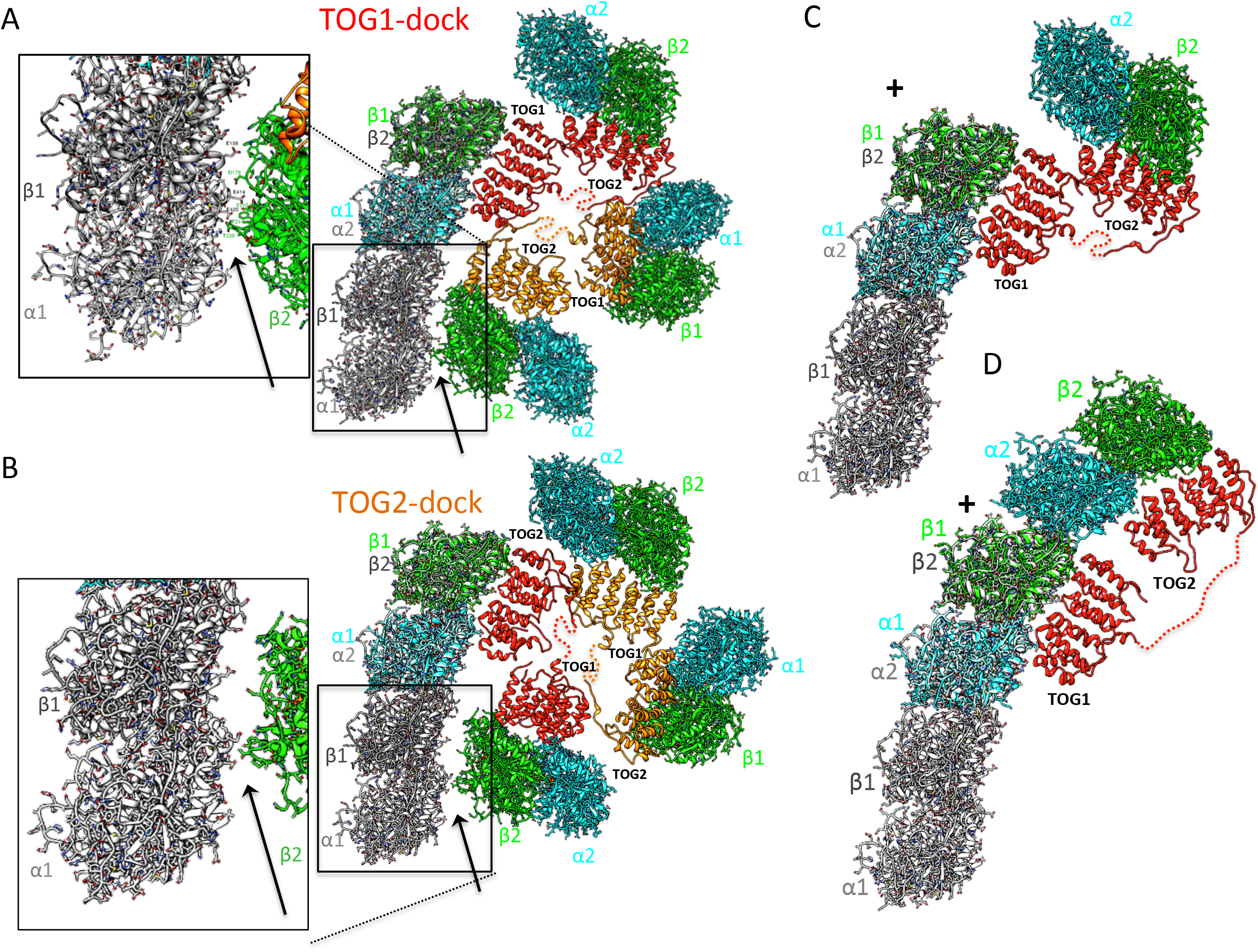
Docking the TOG square and unfurled assembly structures onto protofilament ends at MT plus ends. (**A**) Right, Atomic model for four αβ-tubulin bound TOG square x-ray structure (Figure 2) were docked using αβ-tubulin bound onto TOG1 to the terminal αβ-tubulin in a curved protofilament (PDB ID: 3RYH). Left, magnified view of the zone of steric contact between TOG2-αβ-tubulin in the second subunit and the penultimate αβ-tubulin of the protofilament below the polymerization site. (**B**) Right, Atomic model for four αβ-tubulin bound TOG square x-ray structure (Figure 2) docked using αβ-tubulin bound onto TOG2 to the terminal αβ-tubulin in a curved protofilament (PDB ID: 3RYH). Left, magnified view of the zone shown in A between TOG1-αβ-tubulin in the second subunit and the penultimate αβ-tubulin of the protofilament. Details and overlay images are shown in Figure 7 Supplement 1. (**C**) Docking the isolated TOG1-TOG2 two αβ-tubulin assembly structure (extracted from the dimer structure) onto the terminal αβ-tubulin of the curved protofilament (**D**) docking of the unfurled 1:2:1 unfurled assembly structure (Figure 5) onto the curved protofilament revealing TOG1 to be positioned at the base of the new assembly while TOG2 lies at the outer end of the newly formed MT plus-end.

## Discussion

### A “Polarized Unfurling” model for TOG arrays as MT polymerases

The combination of structural, biochemical analyses suggest a new model for TOG arrays as MT polymerases. We propose a “polar¡zed-unfurl¡ng” MT polymerase model, as summarized in Figure 8 and animated in Video S1, and is supported by docking models for each step (Figure 7) as well as biochemical affinities of TOG1 and TOG2 domains. The polarized unfurling model indicates TOG arrays form two separate states during MT polymerase activity: an αβ-tubulin recruitment complex and plus-end concerted polymerization complex. These two states denoted by the TOG square and unfurled x-ray structures, which we effectively trapped by regulating the polymerization propensity for αβ-tubulin using DRP affinity (Figure 2 and 5). We hypothesize that the association of the αβ-tubulin bound TOG square onto the MT-plus ends, via β-tubulin binding drives the destabilization of these TOG square assemblies and promotes the concerted unfurling of the αβ-tubulins to form new protofilaments.

**Figure 8.**
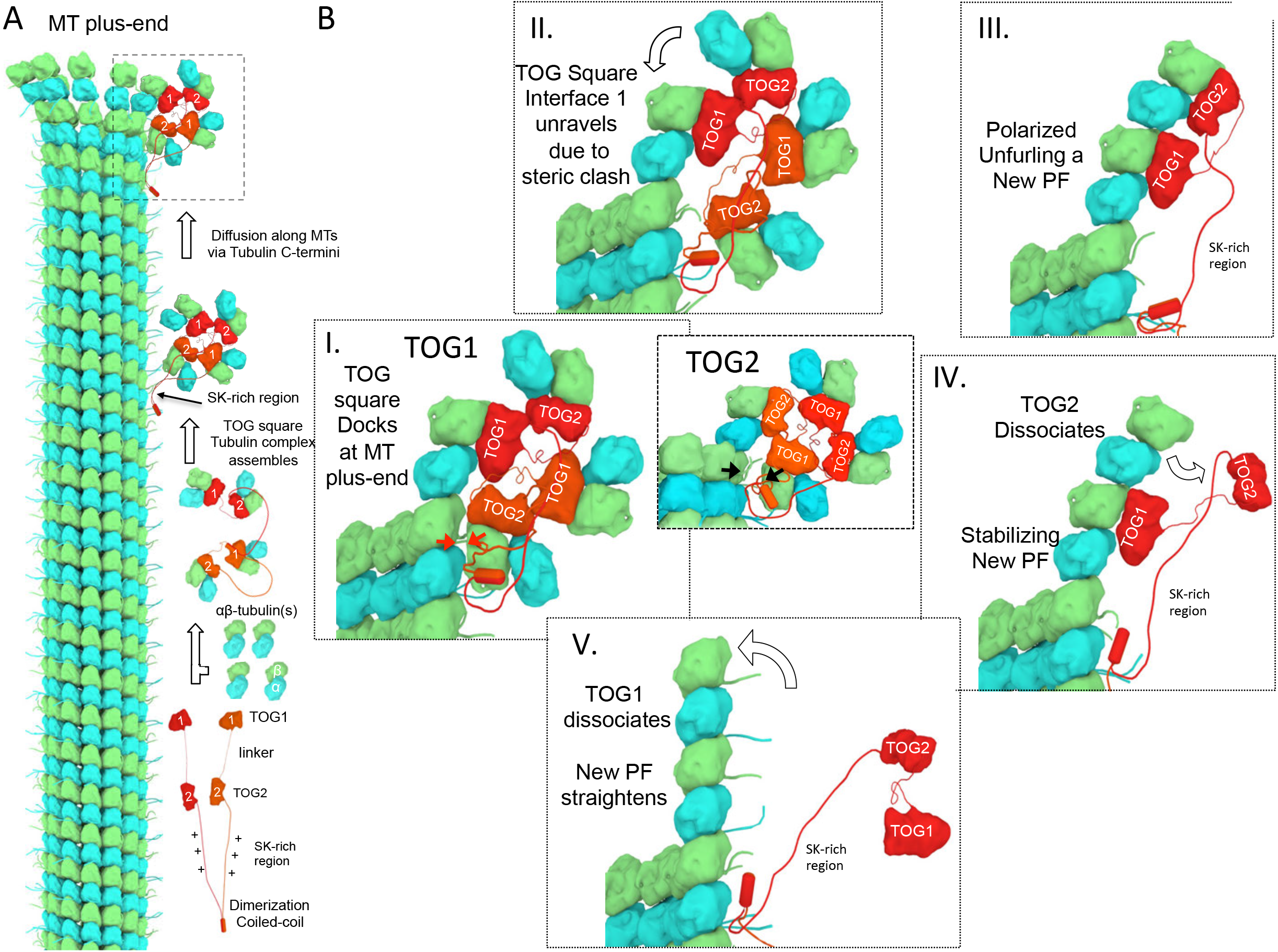
A polarized unfurling model for TOG arrays as MT polymerases. An animation for this model is shown in Video S1. (**A**) Assembly of yeast MT polymerase dimeric TOG1-TOG2 subunits with four αβ-tubulins into a αβ-tubulin: TOG square. TOG squares diffuse along MT lattices modulated by tubulin C-termini interacting with SK-rich regions. (**B**) I. TOG square assemblies orient αβ-tubulins in wheel-shaped assemblies at MT plus ends. II. These assemblies are destabilized upon TOG1-α-tubulin polymerizing onto the exposed β-tubulin at MT plus-ends, releasing TOG1-TOG2 subunits in corner conformation. III. The release of TOG2:αβ-tubulin allows free rotation around TOG1, driving two αβ-tubulins to polymerize. IV. TOG2 dissociates from the newly polymerized αβ-tubulin stabilizing protofilament at plus-end, while TOG1 anchors this αβ-tubulin onto the MT plus end. V. Straightening of this new protofilament leads to the dissociation of TOG1. The rebinding of TOG1-TOG2 subunits to αβ-tubulins reforms the TOG square assembly and restarts the MT polymerase cycle. Atomic views for states I., II., and III are shown in Figure 7 respectively.

We envision the following steps for the polarized unfurling MT polymerase model. First, upon binding four soluble αβ-tubulins, the dimeric TOG1-TOG2 array organizes into TOG square assemblies (Figure 8A). These assemblies place αβ-tubulins in close proximity in a near head-to-tail orientation, while inhibiting their spontaneous polymerization—a feature critical for MT polymerase activity. The TOG square structure reveals that the polarized orientations of recruited αβ-tubulins are due to the asymmetry in each TOG domain:αβ-tubulin interface and the unique head-to-tail assembly of the dimeric TOG1-TOG2 array formed in the TOG square (Figure 2). Positively charged unstructured SK-rich and coiled-coil dimerization regions following the TOG domains may stabilize the TOG square in the wt-Alp14 dimer (Widlund et al., 2011). The propensity of TOG1-TOG2 subunits to form head-to-tail self-assembly is likely strongly enhanced via their dimerization through their c-terminal coiled-coil. In the mammalian proteins such as XMAP215, TOG3-TOG4 likely substitutes for the second subunit of TOG1-TOG2 dimer and may play only a structural role in stabilizing the square conformation (Widlund et al., 2011). Our cysteine crosslinking/mass-spectrometry and mutagenesis studies indicate wt-Alp14-dimer form TOG squares readily in solution, via interfaces 1 and 2. Biochemical and negative stain-EM studies of three mutants (INT1, INT2 and INT12) suggest that interfaces 1 & 2 are responsible for TOG square organization and but they do not influence αβ-tubulin binding. Disabling these interfaces likely leads to poor preorganization of αβ-tubulins and to some degrees of spontaneous αβ-tubulin polymerization while bound to TOG1-TOG2 arrays.

The αβ-tubulin bound TOG square assemblies diffuse along MT lattice, mediated by the SK-rich regions with acidic tubulin C-termini exposed on MT surfaces (Figure 8A) (Al-Bassam et al., 2012; Brouhard et al., 2008). Upon these TOG squares reaching the β-tubulin exposed at MT-plus end protofilament tips, α-tubulin of the TOG1 or TOG2 bound αβ-tubulin may polymerize with β-tubulin exposed at the MT plus-end tip (Figure 8B-I; Figure 2B). The TOG1-αβ-tubulin polymerization is the most likely because of the high occupancy of αβ-tubulin on TOG1 due to its low exchange rate, compared to the lower occupancy of αβ-tubulin on TOG2 due to its rapid exchange rate. Atomic docking models (Figure 8) suggest when a four αβ-tubulin bound TOG square docks onto a protofilament end, steric overlap with the MT protofilament likely destabilizes interface 1 (Figure 8A-I, 8B-I; Figure 7 Supplement 1A). In contrast, TOG squares binding to protofilament ends via TOG2:αβ-tubulin likely would not destabilize at interface 2 due to lack of steric overlap with protofilament end. Thus the asymmetry of the TOG square dimensions may favor TOG square destabilization at interface 1. Thus, we envision this MT plus-end driven TOG square destabilization likely triggers unfurling of a TOG square at MT plus-ends. One or more half-square TOG1-TOG2 subunit corner-like assemblies are then released. The corner-like state of αβ-tubulin bound TOG1-TOG2 subunits are likely unstable (Figure 8B-II). Interface 2 likely acts as a flexible hinge for TOG2 to freely rotate around TOG1 driven by Brownian diffusion, promoting its αβ-tubulin to polymerize catalytically onto the plus-end of the TOG1 bound αβ-tubulin (Figure 8B-III). Our structural studies suggest two TOG1 and TOG2 bound αβ-tubulins polymerize in a single concerted unfurling event, as evidenced by a single elongated complex as seen in the unfurled x-ray structure (Figure 5E, F). This effectively “unfurls” a single curved protofilament from two αβ-tubulins pre-aligned on the TOG1-TOG2 corner of a TOG square assembly. TOG array unfurling activity is directly driven by Brownian diffusion and is captured by αβ-tubulin polymerization through forming inter-dimer polymerization interfaces as shown in our structural studies (Figure 6A, B). The αβ-tubulin inter-dimer interface (1650-Å surface area in a single interface) is fairly sufficient outcompete with TOG square (1930 Å in total for TOG square) reformation.

The unfurled TOG1-TOG2 αβ-tubulin assembly structure reveals that a gradient in αβ-tubulin affinity and exchange rate exists across TOG1 and TOG2 (Figure 1). This gradient is positioned spatially across lower and upper positions of the newly formed protofilament with respect to the MT plus end. The tighter binding affinity of TOG1 for αβ-tubulin likely anchors the TOG array onto the MT plus end, while the rapid exchange of αβ-tubulin by TOG2 likely drives polymerization followed by release. The polarized unfurling model suggests that TOG1 and TOG2 serve specific roles in the MT polymerase activity. This is in contrast to random or reversed orientations that were suggested by other groups (Ayaz et al., 2014; Fox et al., 2014). In a newly unfurled protofilament TOG2 dissociates from upper αβ-tubulin, rapidly, while TOG1 remains tightly bound to the lower αβ-tubulin (Figure 1; Figure 1 Supplement 1). Thus we envision that during MT polymerase cycle that a TOG1-anchored curved protofilament with TOG2 dissociated is a critical intermediate favoring MT polymerization (Figure 8B IV). Protofilament straightening during MT plus-end closure likely induces TOG1 dissociation from the lower αβ-tubulin of the newly polymerized protofilament (Figure 8B-V) (video S1). The studies described in the accompanying manuscript provide in vitro and in vivo evidence for the unique and non-additive roles for TOG1 and TOG2 domains in MT-plus end tracking and MT polymerase activities of Alp14, as supporting the predictions made here (Cook et al., 2018).

The polarized unfurling model explains two observations that were previously not rationalized: 1) fusion of large masses such as GFP protein and his-tag onto the TOG1 N-terminus severely inactivates MT polymerases, activate their MT depolymerization activity, without affecting αβ-tubulin binding (Lechner et al., 2012); Our model shows that N-terminal fusions onto TOG arrays strongly interfere with formation of a TOG square conformation by steric hindering interface 2 assembly. 2) Duplication of TOG1 domains in the yeast does not rescue loss of TOG2 and their sequence positions and length of linkers are critical (Al-Bassam et al., 2006). The TOG square structure suggests that the likely loss of unique interactions that interfaces 1 and 2 form between TOG1 and TOG2 domains in TOG square.

The polarized unfurling model predicts unique roles for the αβ-tubulin recruitment activities by TOG1 and TOG2 domains, and for their assembly into TOG square organization during MT polymerase activity. Here we describe two classes of mutants with defects in recruitment (TOG1M and TOG2M) and TOG square organization (INT1, INT2 and INT12). One prediction is that TOG domains serve distinct and non-additive roles in a concerted MT polymerase mechanism. Our model suggests that TOG1 is essential for anchoring TOG arrays to MT plus ends, while TOG2 is critical for MT polymerase activity. This predicts unique contributions to MT polymerase and MT plus-end tracking activities. A second prediction is that TOG square assembly interfaces are critical for processive MT polymerase at MT plus-ends. In the accompanying manuscript (Cook et al., 2018), we study the two classes of mutants and test predictions of the polarized unfurling MT polymerase model using *in vitro* MT dynamic reconstitution studies and *in vivo* functional studies in *S. pombe* cells.

Other conserved classes of MT regulatory TOG-like array proteins, such as CLASP and Crescerin/CHE-12 may form similar TOG-square like particles and modulate MT dynamics in related mechanisms. For instance, the *S. pombe* CLASP:αβ-tubulin complexes form wheel-like particles with similar dimensions, and promote local MT rescue (Al-Bassam and Chang, 2011; Al-Bassam et al., 2010; Das et al., 2015). Taken together, our data provide a new model for a multi-step MT polymerase mechanism that accelerates αβ-tubulin polymerization.

## Acknowledgement

We thank Dr. Julian Eskin (Brandeis University) for animating the microtubule polymerase mechanism. We thank Advanced Photon Source (APS) and Dr. K. Rajashankar and J. Schuermann, N. Sukumar and D. Neau of the Northeastern Collaborative Access Team (NE-CAT) in using the 24-IDC and IDE beam lines to collect all X-ray diffraction data for our crystallographic studies. We thank Dr. Christopher Fraser for using his Nano-ITC. We thank Dr Rick Mckenney (UC-Davis), and Dr Kevin Corbett (UC-San Diego) for advice and critical reading of this manuscript. JAB and FC are supported by National Institutes of Health GM110283 and GM115185, respectively. sk-Alp14-monomer: αβ-tubulin: DRP, sk-Alp14-monomer-SL:αβ-tubulin: DRP and sk-Alp14-monomer:αβ-tubulin: DRP-ΔN under PDB-ID XXXX, PDB-ID XXXX and XXXX PDB-ID XXXX, respectively.

## Author contribution

JAB, SN and BC designed experiments. BC and SN cloned, expressed and purified Alp14 TOG arrays proteins and carried out biochemical studies. SN determined TOG domain affinities. SN purified and crystallized and optimized TOG array tubulin complexes, determined DRP and DRP-ΔN strategy determined and refined x-ray structures. JAB and SN interpreted structures and built models. JAB, SN and BC designed studies with cysteine mutants and BC carried out studies. JAB carried out cryo-EM image analyses of TOG array complexes. BC carried out studies with inactivated using TIRF microscopy. All Authors wrote, revised the manuscript, created and revised figures.

## Materials and Methods

### Protein expression and purification of Alp14 and sk-Alp14 proteins

The coding regions for MT polymerases from *Saccharomyces cerevisiae* Stu2p (accession: CAA97574.1), *Saccharomyces kluyveri* Stu2p (coding region identified in accession: SKLU-Cont10078), *Schizosaccharomyces pombe* Alp14p (accession: BAA84527.1), and *Chaetomium thermophilum* Stu2 (accession: XP_006692509) were inserted into bacterial expression vectors with a C-terminal his-tag. wt-Alp14-monomers (residues 1-510), wt-Alp14-dimer (residues 1-690), sk-Alp14-monomer (residues 1-550), sk-Alp14-dimer (residues 1-724), Sc-Stu2-dimer (residues 1-746) and Ct-Stu2-dimer (residues 1-719) constructs were generated, including with or without the Ser-Lys rich and coiled-coil dimerization regions. TOG1M and TOG2M mutants were generated via point mutagenesis of Trp23Ala and Arg109Ala to inactivate TOG1 (TOG1M) and Trp300Ala, Lys381Ala to inactivate TOG2 (TOG2M) domains (Al-Bassam et al., 2012). INT12 mutant was generated via gene synthesis (Epoch life sciences) by introducing 15 residue mutations into wt-Alp14-dimer sequence (L206A L208A F275R D276A L277A V278A K320L R359A L39D S40A D42A L437D S440A E478A R479A). The INT1 and INT2 mutants were generated by a PCR swapping strategy of INT12 with wt-Alp14-dimer strategy leading to INT1 with 8-residue mutations (L206A L208A F275R D276A L277A V278A K320L R359A) and INT2 with 7-residue mutations (L39D S40A D42A L437D S440A E478A R479A). Generally, constructs were transformed and expressed in BL21 bacterial strains using the T7 expression system, and were grown at 37°C and induced with 0.5 mM isopropyl thio-β-glycoside at 18 °C overnight. Cells were centrifuged and then lysed using a microfluidizer (Avastin) Extracts were clarified via centrifugation at 18,000 x *g.* Proteins were purified using Ni-IDA (Macherey-Nagel) and/or ion exchange using Hitrap-SP or Hitrap-Q chromatography followed by size exclusion chromatography using a Superdex 200 (30/1000) column (GE Healthcare). DRP was synthesized (Gene Art, Life Technologies), inserted into bacterial expression vectors with a C-terminal 6Xhis tag, and expressed as described above. Proteins were purified using Ni-NTA (Macherey-Nagel) followed by Hitrap Q ion exchange and followed by size exclusion chromatography as described above. Purified proteins were used immediately or frozen in liquid nitrogen for future use.

### Biochemical analyses of Alp14: αβ-tubulin complexes

Soluble porcine αβ-tubulin (10 μM or 20 μM) purified using two GTP-polymerization cycles at high ionic strength as previously described (Castoldi and Popov, 2003) was mixed with 5 μM *S. kluyveri* (sk) or *S. pombe* wt-Alp14-monomer, wt-Alp14-dimer, TOG1M, TOG2M, INT1, INT2, INT12 mutant proteins then diluted 5 fold. To assess αβ-tubulin assembly, The above protein mixtures were analyzed by mixing the above proteins into 0.5-mL volumes and injected them into a Superdex 200 (10/300) size exclusion chromatography (SEC) column equilibrated in 100 mM or 200 mM KCl binding buffer (50 mM HEPES [pH 7.0], 1 mM MgCl_2_, and 1 mM β-mercaptoethanol with 100 mM KCl or 200 mM KCl) using an AKTA purifier system (GE Healthcare). Elution fractions (0.5 mL) were collected and analyzed via sodium dodecyl sulfate polyacrylamide gel electrophoresis (Bio-Rad). The αβ-tubulin- and Alp14-containing bands were quantitated using densitometry to determine the amounts of bound and unbound αβ-tubulin in each SEC fraction. Molecular masses of wt-Alp14-monomer, wt-Alp14-dimer, TOG1M, TOG2M, INT1, INT2 and INT12 proteins, αβ-tubulin, and their complexes were measured using SEC coupled multi-angle light scattering (SEC-MALS). Complexes were separated on Superdex 200 SEC columns (GE Healthcare) while measuring UV absorbance (Agilent 1100-Series HPLC), light scattering (Wyatt Technology miniDAWN TREOS), and refractive index (Wyatt Technology Optilab T-rEX). Concentration-weighted molecular masses for each peak were calculated using ASTRA v. 6 software (Wyatt Technology).

Isothermal titration calorimetery (ITC) was performed using a Nano-ITC (TA Instruments) to determine DRP and DRPΔN affinities for αβ-tubulin. Experiments were performed at 25 °C. Soluble αβ-tubulin, DRP and DRP-ΔN were diluted in 50 mm HEPES buffer, pH 7.3, 100 mM KCl, 1 mM MgCl2 and 50 μM GDP. The sample cell was filled with tubulin (20-40 μM) for every experiment. 135-250 μM of DRP or DRP-ΔN solutions were injected in volumes of 2 or 5 μL in a series of controlled doses into the sample cell. To determine TOG1 and TOG2 affinities for αβ-tubulin with DRP, proteins were diluted in 50 mm HEPES buffer, pH 7.3, 100 or 200 mM KCl and 1 mM MgCl2. 100-250 μM of TOG1 or TOG2 solutions were injected in volumes of 2 or 5 μL in a series of controlled does into the sample cell containing 1:1 molar ratio of αβ-tubulin and DRP (20-40 μM). The results were analyzed the NanoAnalyze software (TA Instruments). Thermodynamic parameters were calculated using the standard thermodynamic equation: *-RTlnK_a_* = *ΔG* = *ΔH-TΔS,* where ΔG, ΔH, and *ΔS* are the changes in free energy, enthalpy, and entropy of binding, respectively, *T* is the absolute temperature, and *R* is the gas constant (1.98 cal mol^−1^ K^−1^).

### Crystallization of Alp14:αβ-tubulin: DRP or DRP-ΔN complexes

Complexes (200 μM) were screened for crystallization using commercial sparse matrix (Qiagen) or homemade screens in 96-well format using a Mosquito robot (TTP Labtech) via the hanging drop method. Cube-shaped crystals (5 μm on each edge) formed for *S. kluyveri* sk-Alp14-monomer:αβ-tubulin: DRP complexes and grew over 4-7 days in 50 mM PIPES, 100 mM MgCl_2_ [pH 7.0], and 10-15% PEG-8000. Larger crystals were grown using micro-seeding (Figure 2-Supplement 2A). To obtain improved X-ray diffraction (see below), we used a sk-Alp14-monomer construct in which non-conserved 270-295 residue linkers was replaced by the shorter linker from the *K. lactis* ortholog sequence (termed sk-Alp14-monomer-SL). Crystals were transferred to 50 mM PIPES, 100 mM MgCl_2_ [pH 7.0], 15% PEG-8000, and 25% glycerol for cryo-protection and flash frozen in liquid nitrogen.

Rectangular crystals of *S. kluyveri* sk-Alp14-monomer:αβ-tubulin: DRP-ΔN complexes formed in 7-10 days under the same conditions described for cube-shaped TOG1-TOG2:αβ-tubulin: DRP crystals. These rectangular crystals exclusively formed using DRPΔN (did not form with DRP), and were obtained using a variety of constructs of monomeric as well as dimeric *S. kluyveri* sk-Alp14-monomer (Table S3). Rectangular sk-Alp14-monomer: αβ-tubulin: DRP-ΔN crystals were treated for cryo-protection and flash frozen as described above.

### X-ray diffraction and structure determination of sk-Alp14:αβ-tubulin assemblies

More than 100 sk-Alp14-monomer:αβ-tubulin: DRP crystals were screened for X-ray diffraction at the Argonne National Laboratory at the Advanced Photon Source microfocus 24-ID-C beamline. Anisotropic X-ray diffraction data were collected for the best cube-shaped crystals in the P2i space group to 4.4 Å resolution in the best dimension, with unit cell dimensions a=216 Å, b=109 Å, and c=280 Å (Figure 2 Supplement 1). The sk-Alp14-monomer-SL:αβ-tubulin-DRP crystals showed improved diffraction and decreased anisotropy to 3.6-Å resolution in an identical P2_1_ unit cell (Table S3). X-ray diffraction data were indexed and scaled using iMOSFLM and treated for anisotropic diffraction using ellipsoidal truncation on the UCLA server (services.mbi.ucla.edu/anisoscale). Phase information was determined using TOG1 (PDB ID:4FFB), TOG2 (PDB ID:4U3J), αβ-tubulin dimer, and DRP (PDB ID:4DRX) models using molecular replacement. Briefly, a truncated poly-alanine TOG domain including only its HEAT repeats was used in the molecular replacement rotation and translation search (Figure 2 Supplement 1B-C). Eight αβ-tubulin and TOG domain solutions were identified based on the Log Likelihood Gain (LLG) values (Figure 2 Supplement 1B-C). After eight cycles of density modification, the electron density map revealed the TOG1 domains exclusively due to the unique C-terminal linker and vertical helix densities (Figure 2 Supplement 1D-E). Density for eight DRP molecules was identified and built. DRP molecules interacted only with their cognate β-tubulin and did not form interfaces with α-tubulin from the neighboring molecule (Figure 2 Supplement 1H). Two 2:4:4 sk-Alp14-monomer:αβ-tubulin: DRP wheel-like models were built and subjected to cycles of rigid-body refinement and model building using the *S. kluyveri* ortholog sequence. Each asymmetric unit contains two wheel-like assemblies (Figure 2 Supplement 1E). TOG1-TOG2 linker residues 270-295 were not observed and were presumed to be disordered (Figure 2 Supplement 1F-H). Density maps from each of the wheel-like models were averaged using non-crystallographic symmetry and then refined using the PHENIX program (Adams et al., 2010). Initially, models were refined using non-crystallographic symmetry (16 fold NCS) restraints and strictly constrained coordinates with group B-factor schemes. In the final stage refinement, the strategy was switched to individual positional and isotropic B-factor with automatic weight optimization. A 4.4-Å sk-Alp14-monomer:αβ-tubulin: DRP structure and 3.6-Å sk-Alp14-monomer-SL:αβ-tubulin:DRP structure are reported; refinement statistics appear in Table S3.

Rectangular crystals formed from sk-Alp14-monomer:αβ-tubulin: DRPΔN diffracted to 3.3-Å resolution at the Argonne National Laboratory at the Advanced Photon Source microfocus APS 24-IDC beamline. X-ray diffraction data were indexed in the P2_1_ space group with unique unit cell dimensions a=115 Å, b=194 Å, and c=149 Å, with two complexes in each unit cell (table S3). Phase information was determined using molecular replacement using the TOG1, TOG2 domains and curved αβ-tubulin as search models (Figure 5 Supplement 1B). TOG1 and TOG2 domains were identified after cycles of density modification as described above. Four αβ-tubulins, four TOG domains, and two DRPΔN models were placed in the unit cell. The identity of TOG domains was determined using the conserved C-terminal linker and jutting helix in the TOG1 domain sequence. A single DRPΔN molecule was identified bound per two αβ-tubulins polymerized complex. Data from each extended assembly were combined using non-crystallographic symmetry (8 fold NCS) and were averaged and refined using the program PHENIX (Adams et al., 2010) (Table S3). The individual positional coordinates and isotropic B-factor were refined with automatic weight optimization in the final stage. A 3.3-Å resolution refined density map is presented in Figure 5 Supplement 1C. Examining data quality of sk-Alp14-monomer:αβ-tubulin: DRP or sk-Alp14-monomer:αβ-tubulin: DRP-ΔN using PHENIX (Adams et al., 2010) indicated the diffraction data contained a small degree of pseudo-merohedral twinning. The twin fractions were adjusted during refinement of both models.

### Cysteine-crosslinking analyses of sk-Alp14 interfaces

Based on the sk-Alp14-monomer:αβ-tubulin: DRP crystal structure, *S. kluyveri* (sk-Alp14) ortholog protein, in its dimer form (residues 1-724), was used to generate crosslinking mutations. Interface 1 residues, which are in close proximity to each other, were mutated to cysteine: Ser180Cys (S180C) and Leu304Cys (L304C), which we termed S180C-L304C. Interface 2 residues, which are in close proximity to each other, were also mutated to cysteine: Ser41Cys (S41C) and Glu518Cys (E518C), which we termed S41C-E518C. The *S. kluyveri* ortholog dimer S180C-L304C mutant and S41C-E518C mutant proteins were purified as described above (Figure 3E). These constructs were either used directly or used to make complexes with αβ-tubulin in a 2:4 (subunit:αβ-tubulin) molar ratio, as described in Figure 1A. These S180C-L304C and S41C E518C mutants, or their αβ-tubulin complexes were then treated using 5 mM Cu-phenanthroline in 50 mM HEPES and 100 mM KCl, pH 7.0 for 5 min, then treated with 5 mM EDTA. These protein mixtures were subjected to sodium dodecyl sulfate polyacrylamide gel electrophoresis under oxidizing conditions.

For LC/MS-MS mass-spectrometry based disulfide peptide mapping, S180C-L304C sk-Alp14 oxidized SDS-PAGE bands were subjected to in-gel proteolysis using either Trypsin or Chymotrypsin. Fragmented peptides were then purified and treated with 5 mM iodioacetamide, which covalently adds 57-Da in mass onto reduced cysteine containing peptides, and does not affect cysteines locked in disulfides. The peptide mixture was then treated with 5 mM Dithiothriatol to reduce disulfides and then treated with 5-Vinyl chloride, which covalently adds 105-Da mass units onto newly reduced cysteine containing peptides. LCMS/MS Mass-spectrometry was performed and the resulting peptides were analyzed. Peptides covering 90% the sk-Alp14 were identified and majority of cysteines were identified. Only two peptides were identified with cysteine-residues that include 105-Da mass units added as described in Figure 4, S6D.

### Negative stain electron microscopy of TOG-array:αβ-tubulin

SEC purified αβ-tubulin complexes of wt-Alp14-dimer, INT1, INT2 and INT12 were placed on glow discharged grids and blot after 30-60 seconds and then stained with multiple washes of 0.1% Uranyl Formate at pH 7. Grids were imaged using low-dose mode at 200 KeV using a DE-20 direct electron detector device (DDD) operating in integration mode.

### Animating the MT polymerase “polarized Unfurling” mechanism

The animation was created using BLENDER 3D-animation software (http://blender.org) as follows: Briefly, surface and ribbon models of PDB coordinates representing the structures, were exported from UCSF-Chimera and were imported into BLENDER, and were then smoothed and optimized to generate cartoon models. Additional protein SK-rich regions and coiled-coil domain structures are unknown structure were thus modeled using sequence length and other information as guidance. The microtubule lattice was modeled based on the tubulin structure (PDB ID 36JF). The dissociation of TOG1 and TOG2 domains from αβ-tubulins were simulated in the animation, based on biochemical studies described in Figure 1 and S1-S3.

---

**Table S1:** The stoichiometry for MT polymerases TOG1-TOG2 binding αβ-tubulin and DARPin (DRP)

| Protein complex | Expected Mass (kDa) | SEC-MALS Measured Mass (kDa) | SEC elution volume (mL) | Apparent Mass (kDa) |
| --- | --- | --- | --- | --- |
| $\alpha\beta$-Tubulin | 100 kDa | 98 ± 0.323* | 12.9 | 104 |
| Alp14-dimer | 150 kDa | 143 ± 1.70 | 10.0 | 675 |
| 1 Alp14-dimer: 1 Tubulin 80 mM KCl | 350 kDa | 387 ± 2.74 | 9.27 | 463 |
| 1 Alp14-dimer: 2 Tubulin 80 mM KCl | 550 kDa | 578 ± 1.41 | 8.98 | 784 |
| 1 Alp14-dimer: 1Tubulin 200 mM KCl | 350 kDa | 304 ± 11.8 | 9.22 | 693 |
| 1 Alp14-dimer: 2 Tubulin 200 mM KCl | 350 kDa | 392 ± 9.66 | 9.67 | 549 |
| 1 Alp14-dimer-TOG1M: 1 Tubulin 80 mM KCl | 350 kDa | 400 ± 7.3 | 10.13 | 433 |
| 1 Alp14-dimer-TOG2M: 2 Tubulin 80 mM KCl | 350 kDa | 382 ± 14 | 9.76 | 524 |
| Alp14-monomer | 62 kDa | 77.8 ± 1.21 | 12.98 | 99 |
| 1 Alp14-monomer: 2 Tubulin 80 mM KCl | 262 kDa | 264 ± 1.31 | 11.17 | 253 |
| 1 Alp14-monomer: 2 Tubulin: 2 DRP 80mM KCl | 298 kDa | 312 ± 2.32 | 10.94 | 285 |
| 1 Alp14-dimer-INT1: 2 Tubulin 80mM KCl | 550 kDa | 533 ± 3.11 | 8.95 | 775 |
| 1 Alp14-dimer-INT: 2 Tubulin 80mM KCl | 550 kDa | 580 ± 3.11 | 8.90 | 770 |
| 1 Alp14-dimer-INT12: 2 Tubulin 80mM KCl | 550 kDa | 540 ± 3.22 | 8.80 | 790 |

\*Standard error is defined based on fitting data across peaks using Astra-software

**Table 2:** Capacities of MT polymerase TOG1-TOG2 to bind αβ-tubulin, influenced by ionic strength

| **Protein Complex**<br>**(Alp14-monomer concentration 4.43 μM)** | **μM tubulin\***<br>**bound** | **μM tubulin**<br>**free** | **TOG1-TOG2**<br>**:$\alpha\beta$-tubulin** |
| --- | --- | --- | --- |
| **<u>wt-Alp14-monomer (2μM)</u>** |  |  |  |
| 1 Alp14-monomer: 1 Tub 100 mM KCl | 4.57 ± 0.08 | 0.43 ± 0.08 | 1.04 |
| 1 Alp14-monomer: 2 Tub 100 mM KCl | 8.82 ± 0.16 | 1.18 ± 0.16 | 2 |
| 1 Alp14-monomer: 1 Tub 200 mM KCl | 3.37 ± 0.27 | 1.63 ± 0.27 | 0.76 |
| 1 Alp14-monomer: 2 Tub 200 mM KCl | 5.97 ± 0.12 | 4.03 ± 0.12 | 1.35 |
| 1 Alp14-monomer:1 Tub:1 DRP 100 mM KCl | 4.33 ± 0.13 | 0.67 ± 0.13 | 0.98 |
| 1 Alp14-monomer:2 Tub:2 DRP 100 mM KCl | 7.23 ± 0.01 | 2.77 ± 0.01 | 1.64 |
| 1 Alp14-monomer:1 Tub:1 DRP 200 mM KCl | 4.13 ± 0.09 | 0.87 ± 0.09 | 0.94 |
| 1 Alp14-monomer: 2 Tub 2 DRP 200 mM KCl | 5.67 ± 0.03 | 4.34 ± 0.03 | 1.28 |
| **<u>wt-Alp14-dimer (1μM)</u>** |  |  |  |
| 1 Alp14-dimer: 1 Tub 100 mM KCl | 4.31 ± 0.07 | 0.69 ± 0.07 | 0.98 |
| 1 Alp14-dimer: 2 Tub 100 mM KCl | 7.11 ± 0.14 | 2.89 ± 0.14 | 1.61 |
| 1 Alp14-dimer: 1 Tub 200 mM KCl | 3.44 ± 0.37 | 1.56 ± 0.37 | 0.78 |
| 1 Alp14-dimer: 2 Tub 200 mM KCl | 5.46 ± 0.34 | 4.55 ± 0.34 | 1.24 |
| 1 Alp14-dimer: 1 Tub: 2 DRP 100 mM KCl | 4.23 ± 0.02 | 0.78 ± 0.02 | 0.96 |
| 1 Alp14-dimer: 1 Tub: 2 DRP 100 mM KCl | 8.33 ± 0.09 | 1.67 ± 0.09 | 1.89 |
| 1 Alp14-dimer: 1 Tub: 1 DRP 200 mM KCl | 4.17 ± 0.29 | 0.83 ± 0.29 | 0.95 |
| 1 Alp14-dimer: 2 Tub: 2 DRP 200 mM KCl | 6.02 ± 0.17 | 3.99 ± 0.17 | 1.36 |
| **<u>TOG square Interface mutants (1μM)</u>** |  |  |  |
| 1 INT1: 2 Tub 100mM KCl | 8.34 ± 0.07 | 1.66 ± 0.07 | 1.74 |
| 1 INT1: 2 Tub 200mM KCl | 5.67 ± 0.04 | 4.33 ± 0.04 | 1.18 |
| 1 INT2: 2 Tub 100mM KCl | 9.52 ± 0.3 | 0.48 ± 0.2 | 2.02 |
| 1 INT2: 2 Tub 200mM KCl | 6.82 ± 0.2 | 3.1 ± 0.1 | 1.45 |
| 1 INT12: 2 Tub 100mM KCl | 8.44 ± 0.05 | 1.56 ± 0.05 | 1.86 |
| 1 INT12: 2 Tub 200mM KCl | 6.85 ± 0.04 | 3.15 ± 0.04 | 1.46 |
| **<u>Inactivated TOG mutants (1μM)</u>** |  |  |  |
| 1 TOG1M: 1 Tub 100 mM KCl | 4.04 ± 0.07 | 0.97 ± 0.07 | 0.92 |
| 1 TOG1M: 2 Tub 100 mM KCl | 4.94 ± 0.06 | 5.06 ± 0.06 | 1.12 |
| 1 TOG1M: 1 Tub 200 mM KCl | 1.66 ± 0.25 | 3.34 ± 0.25 | 0.38 |
| 1 TOG1M: 2 Tub 200 mM KCl | 2.23 ± 0.1 | 7.77 ± 0.1 | 0.51 |
| 1 TOG2M: 1 Tub 100 mM KCl | 3.53 ± 0.04 | 1.48 ± 0.04 | 0.8 |
| 1 TOG2M: 2 Tub 100 mM KCl | 6.24 ± 0.10 | 3.76 ± 0.1 | 1.41 |
| 1 TOG2M: 1 Tub 200 mM KCl | 3.15 ± 0.24 | 1.85 ± 0.24 | 0.72 |
| 1 TOG2M: 2 Tub 200 mM KCl | 4.28 ± 0.18 | 5.72 ± 0.18 | 0.97 |

\*Standard error is defined based on combined data from duplicated SEC runs

**Table S3:**
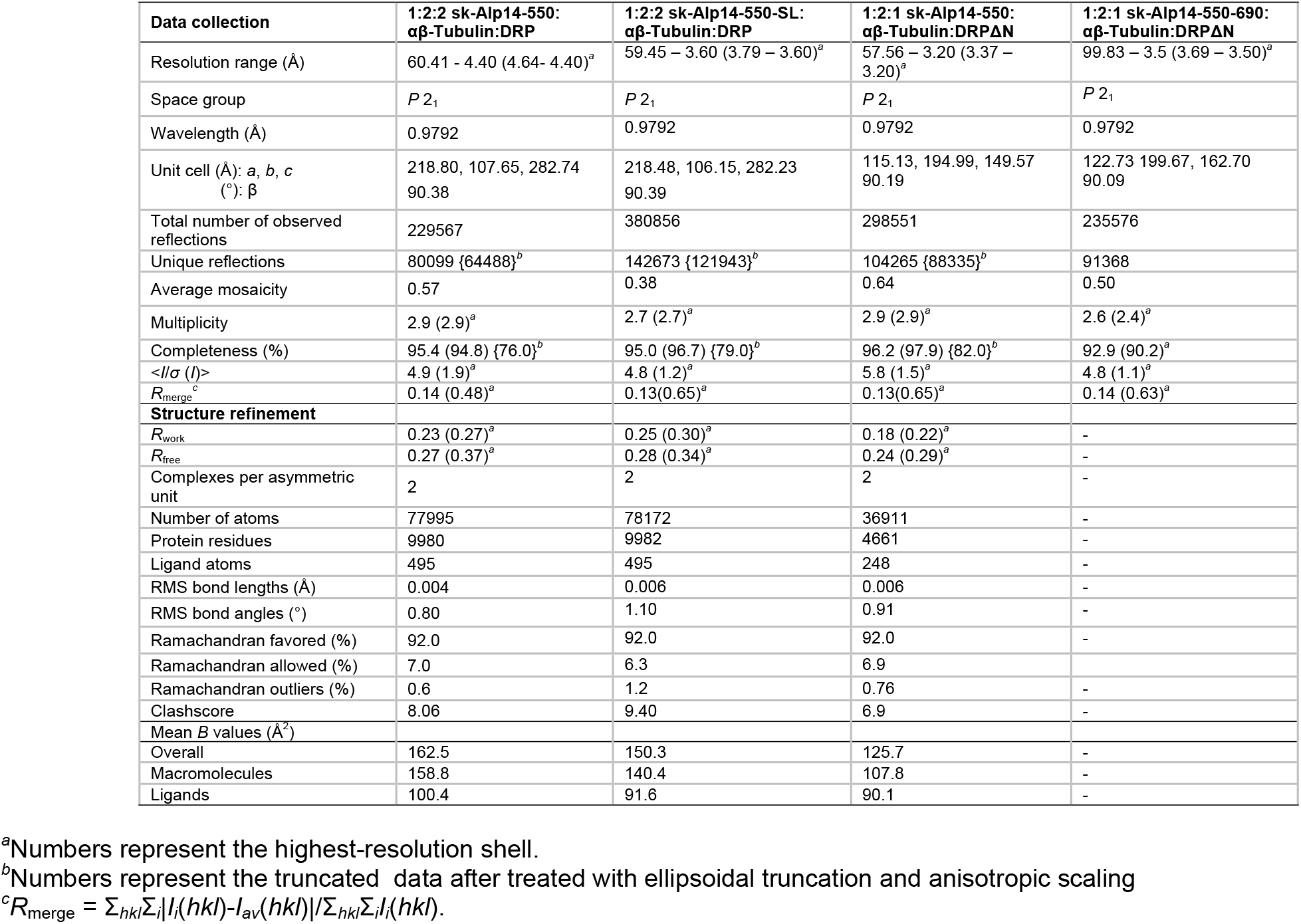
X-ray Crystallographic and Refinement statistics of MT polymerase:αβ-tubulin:DRP

| Data collection | 1:2:2 sk-Alp14-550:<br>$\alpha\beta$-Tubulin:DRP | 1:2:2 sk-Alp14-550-SL:<br>$\alpha\beta$-Tubulin:DRP | 1:2:1 sk-Alp14-550:<br>$\alpha\beta$-Tubulin:DRPΔN | 1:2:1 sk-Alp14-550-690:<br>$\alpha\beta$-Tubulin:DRPΔN |
| --- | --- | --- | --- | --- |
| Resolution range (Å) | 60.41 - 4.40 (4.64- 4.40)<sup>a</sup> | 59.45 - 3.60 (3.79 - 3.60)<sup>a</sup> | 57.56 - 3.20 (3.37 - 3.20)<sup>a</sup> | 99.83 - 3.5 (3.69 - 3.50)<sup>a</sup> |
| Space group | *P* 2<sub>1</sub> | *P* 2<sub>1</sub> | *P* 2<sub>1</sub> | *P* 2<sub>1</sub> |
| Wavelength (Å) | 0.9792 | 0.9792 | 0.9792 | 0.9792 |
| Unit cell (Å): *a*, *b*, *c*<br>(°): $\beta$ | 218.80, 107.65, 282.74<br>90.38 | 218.48, 106.15, 282.23<br>90.39 | 115.13, 194.99, 149.57<br>90.19 | 122.73 199.67, 162.70<br>90.09 |
| Total number of observed reflections | 229567 | 380856 | 298551 | 235576 |
| Unique reflections | 80099 {64488}<sup>b</sup> | 142673 {121943}<sup>b</sup> | 104265 {88335}<sup>b</sup> | 91368 |
| Average mosaicity | 0.57 | 0.38 | 0.64 | 0.50 |
| Multiplicity | 2.9 (2.9)<sup>a</sup> | 2.7 (2.7)<sup>a</sup> | 2.9 (2.9)<sup>a</sup> | 2.6 (2.4)<sup>a</sup> |
| Completeness (%) | 95.4 (94.8) {76.0}<sup>b</sup> | 95.0 (96.7) {79.0}<sup>b</sup> | 96.2 (97.9) {82.0}<sup>b</sup> | 92.9 (90.2)<sup>a</sup> |
| $\langle I/\sigma(I) \rangle$ | 4.9 (1.9)<sup>a</sup> | 4.8 (1.2)<sup>a</sup> | 5.8 (1.5)<sup>a</sup> | 4.8 (1.1)<sup>a</sup> |
| $R_{\text{merge}}^c$ | 0.14 (0.48)<sup>a</sup> | 0.13(0.65)<sup>a</sup> | 0.13(0.65)<sup>a</sup> | 0.14 (0.63)<sup>a</sup> |
| **Structure refinement** |  |  |  |  |
| $R_{\text{work}}$ | 0.23 (0.27)<sup>a</sup> | 0.25 (0.30)<sup>a</sup> | 0.18 (0.22)<sup>a</sup> | - |
| $R_{\text{free}}$ | 0.27 (0.37)<sup>a</sup> | 0.28 (0.34)<sup>a</sup> | 0.24 (0.29)<sup>a</sup> | - |
| Complexes per asymmetric unit | 2 | 2 | 2 | - |
| Number of atoms | 77995 | 78172 | 36911 | - |
| Protein residues | 9980 | 9982 | 4661 | - |
| Ligand atoms | 495 | 495 | 248 | - |
| RMS bond lengths (Å) | 0.004 | 0.006 | 0.006 | - |
| RMS bond angles (°) | 0.80 | 1.10 | 0.91 | - |
| Ramachandran favored (%) | 92.0 | 92.0 | 92.0 | - |
| Ramachandran allowed (%) | 7.0 | 6.3 | 6.9 | - |
| Ramachandran outliers (%) | 0.6 | 1.2 | 0.76 | - |
| Clashscore | 8.06 | 9.40 | 6.9 | - |
| Mean *B* values (Å<sup>2</sup>) |  |  |  |  |
| Overall | 162.5 | 150.3 | 125.7 | - |
| Macromolecules | 158.8 | 140.4 | 107.8 | - |
| Ligands | 100.4 | 91.6 | 90.1 | - |

<sup>a</sup>Numbers represent the highest-resolution shell.<sup>b</sup>Numbers represent the truncated data after treated with ellipsoidal truncation and anisotropic scaling<sup>c</sup> $R_{\text{merge}} = \sum_{hkl} \sum_i |I_i(hkl) - I_{\text{av}}(hkl)| / \sum_{hkl} \sum_i I_i(hkl)$ .

**Table S4:** Buried Surface Area between αβ-tubulin dimer and TOG domains or DRP

| Interface | 1:2:2: sk-Alp14-monomer:<br>$\alpha\beta$-Tubulin: DRP ($\text{\AA}^2$)* | 1:2:2: sk-Alp14-monomer-SL:<br>$\alpha\beta$-Tubulin: DRP ($\text{\AA}^2$) | 1:2:1: sk-Alp14-monomer:<br>$\alpha\beta$-Tubulin: DRP-ΔN<br>($\text{\AA}^2$) |
| --- | --- | --- | --- |
| TOG1 and TOG2-interface 2 | 273 ± 35 | 257 ± 42 | - |
| TOG1-TOG2 dimer-interface 1 | 661 ± 27 | 702 ± 18 | - |
| $\alpha\beta$-tubulin and TOG1 | 752 ± 12 | 786 ± 42 | 804 ± 24 |
| $\alpha\beta$-tubulin and TOG2 | 916 ± 23 | 890 ± 22 | 863 ± 12 |
| β-tubulin and DRP or DRP-ΔN | 842 ± 68 | 873 ± 21 | 846 ± 20 |
| α-tubulin and DRP<br>(inter-$\alpha\beta$-tubulin subunit) | 108 ± 47 | 81 ± 31 | - |

\* Interface areas were determined by a single buried surface, and averaged among each non-crystallographic unit in the structure

**Table S5:** Intra and inter dimer curvature angles (°) of αβ-Tubulins observed in structures

| | PDB ID | Intra dimer ($\alpha 1\beta 1$)<br>angle ($^\circ$) | Inter dimer<br>($\beta 1\alpha 2$) angle ($^\circ$) |
| --- | --- | --- | --- |
| Stathmin:RB3 complex with GTP | 3RYH | 9.2 | 10.3 |
| Stathmin:RB3 complex with GDP | 1SA0 | 13.0 | 13.5 |
| $\alpha\beta$-tubulin:TOG1 complex with GTP | 4FFB | 13 | - |
| $\alpha\beta$-Tubulin:TOG2 complex with GTP | 4U3J | 13 | - |
| $\alpha\beta$-Tubulin:DRP complex with GDP | 4DRX | 11.0 | - |
| Sk-Alp14-monomer:$\alpha\beta$-Tubulin:DRP wheel with GDP | Current | 11.3 | - |
| Sk-Alp14-monomer:$\alpha\beta$-Tubulin:DRP-ΔN with GDP | Current | 11.2 | 16.4 |

\* Curvature of $\alpha\beta$ -tubulin interface were determined as previously described by Rice and Brouhard 2015.

## References

Adams, P.D., Afonine, P.V., Bunkoczi, G., Chen, V.B., Davis, I.W., Echols, N., Headd, J.J., Hung, L.W., Kapral, G.J., Grosse-Kunstleve, R.W., et al. (2010). PHENIX: a comprehensive Python-based system for macromolecular structure solution. Acta Crystallogr D Biol Crystallogr 66, 213–221.

Akhmanova, A., and Steinmetz, M.O. (2008). Tracking the ends: a dynamic protein network controls the fate of microtubule tips. Nat Rev Mol Cell Biol 9, 309–322.

Akhmanova, A., and Steinmetz, M.O. (2011). Microtubule End Binding: EBs Sense the Guanine Nucleotide State. Curr Biol 21, R283–285.

Akhmanova, A., and Steinmetz, M.O. (2015). Control of microtubule organization and dynamics: two ends in the limelight. Nat Rev Mol Cell Biol.

Al-Bassam, J. (2014). Reconstituting dynamic microtubule polymerization regulation by TOG domain proteins. Methods Enzymol 540, 131–148.

Al-Bassam, J., and Chang, F. (2011). Regulation of microtubule dynamics by TOG-domain proteins XMAP215/Dis1 and CLASP. Trends Cell Biol 21, 604–614.

Al-Bassam, J., Kim, H., Brouhard, G., van Oijen, A., Harrison, S.C., and Chang, F. (2010). CLASP promotes microtubule rescue by recruiting tubulin dimers to the microtubule. Dev Cell 19, 245–258.

Al-Bassam, J., Kim, H., Flor-Parra, I., Lal, N., Velji, H., and Chang, F. (2012). Fission yeast Alp14 is a dose-dependent plus end-tracking microtubule polymerase. Mol Biol Cell 23, 2878–2890.

Al-Bassam, J., Larsen, N.A., Hyman, A.A., and Harrison, S.C. (2007). Crystal structure of a TOG domain: conserved features of XMAP215/Dis1-family TOG domains and implications for tubulin binding. Structure 15, 355–362.

Al-Bassam, J., van Breugel, M., Harrison, S.C., and Hyman, A. (2006). Stu2p binds tubulin and undergoes an open-to-closed conformational change. J Cell Biol 172, 1009–1022.

Alushin, G.M., Lander, G.C., Kellogg, E.H., Zhang, R., Baker, D., and Nogales, E. (2014). High-resolution microtubule structures reveal the structural transitions in alphabeta-tubulin upon GTP hydrolysis. Cell 157, 1117–1129.

Ayaz, P., Munyoki, S., Geyer, E.A., Piedra, F.A., Vu, E.S., Bromberg, R., Otwinowski, Z., Grishin, N.V., Brautigam, C.A., and Rice, L.M. (2014). A tethered delivery mechanism explains the catalytic action of a microtubule polymerase. Elife 3, e03069.

Ayaz, P., Ye, X., Huddleston, P., Brautigam, C.A., and Rice, L.M. (2012). A TOG:alphabeta-tubulin complex structure reveals conformation-based mechanisms for a microtubule polymerase. Science 337, 857–860.

Bieling, P., Telley, I.A., Hentrich, C., Piehler, J., and Surrey, T. (2010). Fluorescence microscopy assays on chemically functionalized surfaces for quantitative imaging of microtubule, motor, and +TIP dynamics. Methods Cell Biol 95, 555–580.

Brouhard, G.J., and Rice, L.M. (2014). The contribution of alphabeta-tubulin curvature to microtubule dynamics. J Cell Biol 207, 323–334.

Brouhard, G.J., Stear, J.H., Noetzel, T.L., Al-Bassam, J., Kinoshita, K., Harrison, S.C., Howard, J., and Hyman, A.A. (2008). XMAP215 is a processive microtubule polymerase. Cell 132, 79–88.

Castoldi, M., and Popov, A.V. (2003). Purification of brain tubulin through two cycles of polymerization-depolymerization in a high-molarity buffer. Protein Expr Purif 32, 83–88.

Cook B., Chang. F., Flor-Parra I., Al-Bassam J., (2018) Roles tubulin recruitment and self-organization by Tumor overexpressed gene (TOG) domains array in Microtubule plus-end tracking and polymerase mechanisms. co-submited

Cullen, C.F., Deak, P., Glover, D.M., and Ohkura, H. (1999). mini spindles: A gene encoding a conserved microtubule-associated protein required for the integrity of the mitotic spindle in Drosophila. J Cell Biol 146, 1005–1018.

Das, A., Dickinson, D.J., Wood, C.C., Goldstein, B., and Slep, K.C. (2015). Crescerin uses a TOG domain array to regulate microtubules in the primary cilium. Mol Biol Cell 26, 4248–4264.

Fox, J.C., Howard, A.E., Currie, J.D., Rogers, S.L., and Slep, K.C. (2014). The XMAP215 family drives microtubule polymerization using a structurally diverse TOG array. Mol Biol Cell 25, 2375–2392.

Howard, A.E., Fox, J.C., and Slep, K.C. (2015). Drosophila melanogaster mini spindles TOG3 utilizes unique structural elements to promote domain stability and maintain a TOG1- and TOG2-like tubulin-binding surface. J Biol Chem 290, 10149–10162.

Lechner, B., Rashbrooke, M.C., Collings, D.A., Eng, R.C., Kawamura, E., Whittington, A.T., and Wasteneys, G.O. (2012). The N-terminal TOG domain of Arabidopsis MOR1 modulates affinity for microtubule polymers. J Cell Sci 125, 4812–4821.

Maurer, S.P., Cade, N.I., Bohner, G., Gustafsson, N., Boutant, E., and Surrey, T. (2014). EB1 accelerates two conformational transitions important for microtubule maturation and dynamics. Curr Biol 24, 372–384.

Miller, M.P., Asbury, C.L., and Biggins, S. (2016). A TOG Protein Confers Tension Sensitivity to Kinetochore-Microtubule Attachments. Cell 165, 1428–1439.

Nawrotek, A., Knossow, M., and Gigant, B. (2011). The determinants that govern microtubule assembly from the atomic structure of GTP-tubulin. J Mol Biol 412, 35–42.

Pecqueur, L., Duellberg, C., Dreier, B., Jiang, Q., Wang, C., Pluckthun, A., Surrey, T., Gigant, B., and Knossow, M. (2012). A designed ankyrin repeat protein selected to bind to tubulin caps the microtubule plus end. Proc Natl Acad Sci U S A 109, 12011–12016.

Reber, S.B., Baumgart, J., Widlund, P.O., Pozniakovsky, A., Howard, J., Hyman, A.A., and Julicher, F. (2013). XMAP215 activity sets spindle length by controlling the total mass of spindle microtubules. Nat Cell Biol 15, 1116–1122.

Slep, K.C., and Vale, R.D. (2007). Structural basis of microtubule plus end tracking by XMAP215, CLIP-170, and EB1. Mol Cell 27, 976–991.

Tanaka, K., Mukae, N., Dewar, H., van Breugel, M., James, E.K., Prescott, A.R., Antony, C., and Tanaka, T.U. (2005). Molecular mechanisms of kinetochore capture by spindle microtubules. Nature 434, 987–994.

Wang, P.J., and Huffaker, T.C. (1997). Stu2p: A microtubule-binding protein that is an essential component of the yeast spindle pole body. J Cell Biol 139, 1271–1280.

Widlund, P.O., Stear, J.H., Pozniakovsky, A., Zanic, M., Reber, S., Brouhard, G.J., Hyman, A.A., and Howard, J. (2011). XMAP215 polymerase activity is built by combining multiple tubulin-binding TOG domains and a basic lattice-binding region. Proc Natl Acad Sci U S A 108, 2741–2746.

